# Food and water uptake are regulated by distinct central amygdala circuits revealed using intersectional genetics

**DOI:** 10.1101/2024.06.22.600182

**Authors:** Federica Fermani, Simon Chang, Christian Peters, Louise Gaitanos, Pilar L. Alcala Morales, Charu Ramakrishnan, Karl Deisseroth, Rüdiger Klein

**Author notes:** Correspondence (R.K.).

## Abstract

The central amygdala (CeA) plays a crucial role in defensive and appetitive behaviours. It contains genetically defined GABAergic neuron subpopulations distributed over three anatomical subregions, capsular (CeC), lateral (CeL), and medial (CeM). The roles that these molecularly- and anatomically-defined CeA neurons play in appetitive behavior remain unclear. Using intersectional genetics, we found that neurons driving food or water consumption are confined to the CeM. Separate CeM subpopulations exist for water only versus water or food consumption. *In vivo* calcium imaging revealed that CeM^Htr2a^ neurons promoting feeding are responsive towards appetitive cues with little regard for their physical attributes. CeM^Sst^ neurons involved in drinking are sensitive to the physical properties of salient stimuli. Both CeM subtypes receive inhibitory input from CeL and send projections to the parabrachial nucleus to promote appetitive behavior. These results suggest that distinct CeM microcircuits evaluate liquid and solid appetitive stimuli to drive the appropriate behavioral responses.

## Introduction

The central amygdala (CeA), is a brain hub for emotion and motivation that rapidly integrates salient environmental and internal stimuli to generate appropriate behavioral responses ^1–4^. While traditionally thought to be linked solely to defensive behavior, emerging evidence suggests that specific CeA neuronal populations play key roles in reward and consummatory behaviors, such as feeding ^5, 6^ and drinking ^7^. Since food and water are intrinsically rewarding ^8, 9^, appetitive CeA neurons are positive valence neurons and the animals seek to promote the activation of these neurons. *In vivo* recordings have shown that appetitive CeA neurons respond to appetitive stimuli, such as food and water, as well as to cues predictive of these stimuli ^1, 2, 5, 10–12^. *In vivo* single cell calcium imaging experiments revealed marked heterogeneity in specific appetitive subpopulations, with some appetitive neurons becoming selectively activated by appetitive cues, with others becoming inhibited or showing dynamic responses to multiple stimuli ^5, 12, 13^. In addition to environmental stimuli, appetitive CeA neurons are activated by internal signals, such as the hunger hormone ghrelin ^13^. Despite recent progress, the circuit mechanisms in the central amygdala that process appetitive signals such as food and water are still not well understood ^2, 14–17^.

The CeA consists exclusively of γ-aminobutyric acid-releasing (GABAergic) neurons that can be divided into subpopulations based on their molecular, morphological, and functional properties. They are organized in inhibitory microcircuits in three subregions of the CeA: the central capsular (CeC), lateral (CeL), and medial subregions (CeM). For defensive behavior, the information flow through the subregions has been partially characterized: CeC and CeL subregions are the primary targets for sensory inputs that are processed and passed on to the CeM from where major projections go to hindbrain autonomic and motor control areas ^2, 18, 19^. For appetitive behavior, the information flow is less clear. New perspectives have been suggested, with the CeL also forming long-range efferent projections bypassing the CeM, and the CeM receiving direct inputs bypassing the CeL ^20, 21^.

The functional characterization of molecularly-defined appetitive CeA subpopulations has been the subject of recent intense research. It has mostly relied on the manipulation of single Cre driver lines and expression analysis of selected markers ^5, 7, 22–24^. The more recent characterization of CeA subpopulations by scRNAseq analysis revealed that several of these Cre drivers are expressed in more than one cell cluster and, importantly, across CeA subdivisions ^13, 21, 25^. The Sst-Cre driver has been extensively used to demonstrate that CeA neurons expressing the neuropeptide somatostatin, control both appetitive and aversive behaviors ^7, 11, 12, 26, 27^. *In vivo* calcium imaging suggested that Sst+ CeA neurons help discriminate between stimuli with different physical properties and participate in the evaluation of salient events during reward learning ^12^ . CeA^Sst^ neurons can be found in CeL and CeM subregions and partially overlap with neurons expressing neurotensin (Nts) ^7, 28^ which show a similar distribution pattern and promote consumption of palatable fluids ^23^. The anatomical position of appetitive CeA^Sst^ and CeA^Nts^ neurons has not been fully worked out.

In previous work, we have shown that CeA neurons expressing the Htr2a-Cre driver promote food consumption through a positive valence signal ^5^. CeA^Htr2a^ neurons are activated by fasting, the hunger hormone ghrelin, and the presence of food ^13^. Single cell transcriptomics revealed that CeA^Htr2a^ neurons are located in the CeL and CeM and it is currently unclear which of these populations promotes food consumption. A large fraction of Htr2a-Cre-expressing CeA neurons overlaps with neurons expressing prepronociceptin (Pnoc) ^13^. Similar to CeA^Htr2a^ neurons, CeA^Pnoc^ neurons promote palatable food consumption ^22^, but the exact anatomical location of Pnoc-positive appetitive neurons remains unclear. CeA^Htr2a^ neurons located in the CeL overlap with the Sst-positive population, whereas CeM^Htr2a^ and CeM^Sst^ neurons are largely separate populations ^13^.

To better define the CeA appetitive microcircuits, we employed combinatorial recombinase-dependent targeting ^29^ using available Cre lines in combination with a novel transgenic line that expresses the Flp recombinase specifically in neurons of the CeL subregion. This approach, in combination with INTRSECT Boolean vectors ^30, 31^, allowed for functional characterization of four spatially and molecularly distinct CeA subpopulations, CeL^Sst^, CeM^Sst^, CeL^Htr2a^, and CeM^Htr2a^ neurons. The results revealed a new organization of appetitive information flow through the CeA. Unexpectedly, our results indicate that neurons driving food or water consumption are confined to the CeM. Separate CeM subpopulations exist for water only (CeM^Sst^), and water or food consumption (CeM^Htr2a^). *In vivo* calcium imaging revealed that CeM^Htr2a^ neurons promoting feeding or drinking are highly responsive towards appetitive cues with little regard for their physical attributes. CeM^Sst^ neurons involved exclusively in drinking are sensitive to the physical properties of salient stimuli. Both CeM subtypes are controlled by inhibitory signals from the CeL and, in turn, form long-range inhibitory projections to the PBN to promote water or food consumption. These results suggest that distinct CeM microcircuits evaluate liquid and solid appetitive stimuli to drive the appropriate behavioural responses.

## RESULTS

### A new FlpoER transgenic line for intersectionally targeting central amygdala neurons

To manipulate CeL versus CeM neurons with an intersectional genetics strategy, we generated a Wfs1-FlpoER mouse line that expresses the tamoxifen-inducible optimized FlpoER recombinase under the control of the Wolframin1 (Wfs1) promoter (Fig. 1a). Wfs1 was previously shown to mark the majority of cells in the CeL ^32, 33^ and we confirmed that Wfs1 immunoreactivity was enriched in the CeL (Fig. 1b). Using Htr2a-Cre and Sst-Cre driven tdTomato reporter mice, we found that nearly 90% of Htr2a-Cre and Sst-Cre positive cells expressed Wfs1 in the CeL, and a large fraction of Wfs1 immunoreactive cells encompassed Htr2a-Cre and Sst-Cre positive cells (Extended Data Fig. 1a-j). To validate the accuracy of FlpoER expression, we bred Wfs1-FlpoER mice with a FPDI reporter line which expresses mCherry after Flp-mediated recombination ^34^. After three tamoxifen injections, 56% of cells in CeL that were immunopositive for endogenous Wfs1 expressed FlpoER (mCherry), and, importantly for our intersectional targeting strategy, mCherry+ cells were 17-times more abundant in CeL than CeM (Extended Data Fig. 1k-o).

**Figure 1.**
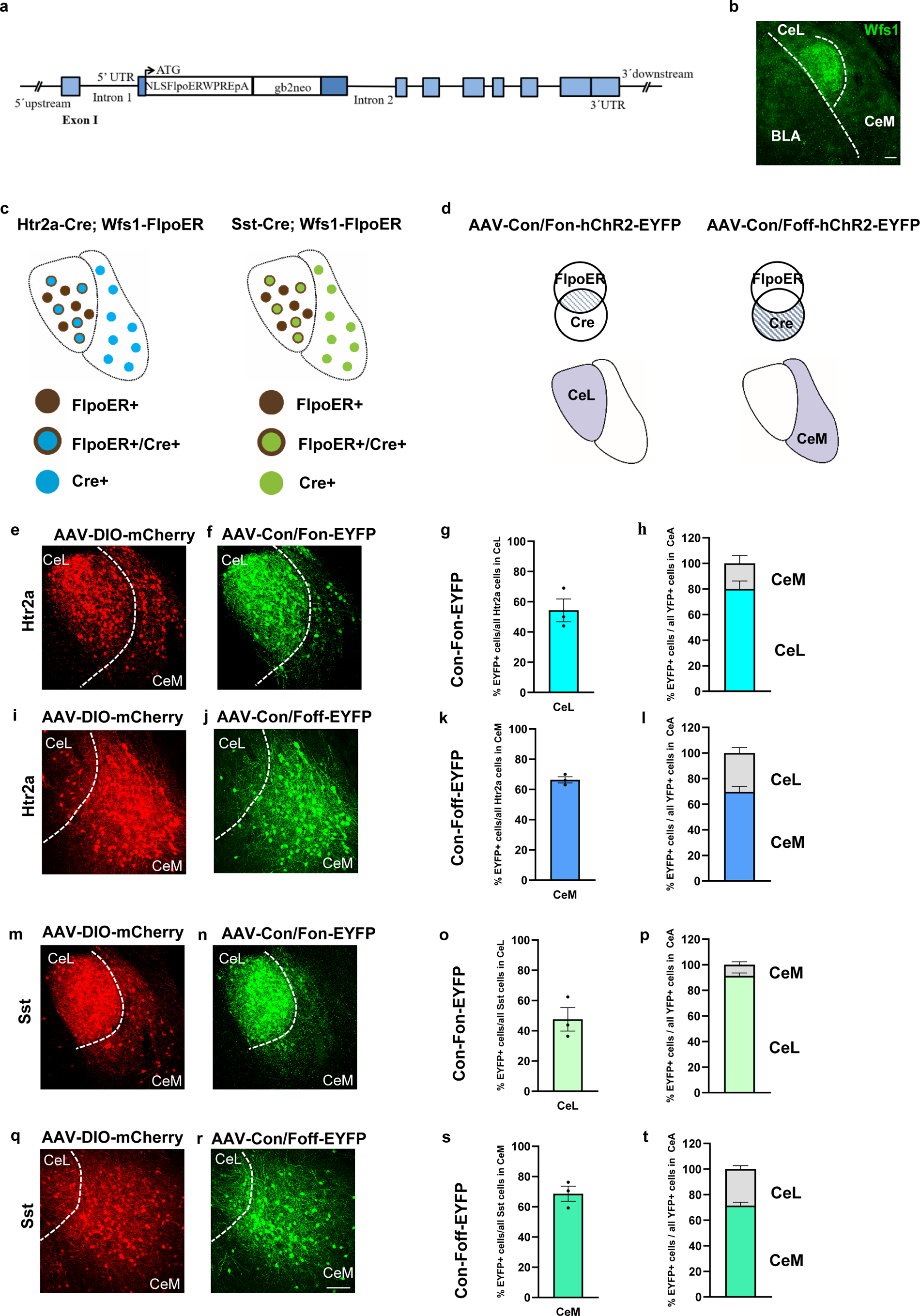
Characterization of the CeL-specific Wfs1-FlpoER driver line. **a.** Scheme of the Wfs1-FlpoER transgene. The Wfs1 BAC clone was targeted with the optimized Flp-estrogen receptor fusion protein (FlpoER) tagged with a nuclear localization sequence (NLS) and linked to the woodchuck hepatitis post-transcriptional regulatory element (WPRE) and bovine growth hormone polyadenylation signal (pA), and a gb2 prokaryotic promoter driven Neo cassette. **b.** Example image illustrating endogenous Wfs1 immunoreactivity in the CeL. **c.** Schemes of the intersectional strategy used with Htr2a-Cre;Wfs1-FlpoER and Sst-Cre;Wfs1-FlpoER mice. Cells in the CeL can be either FlpoER positive or FlpoER and Cre double positive. Cells in the CeM are only Cre-positive. **d.** Schemes of CeA subregional expression of INTRSECT viruses. Con/Fon viruses express in CeL cells that are double positive for Cre and FlpoER, while Con/Foff viruses express in CeM cells that are Cre-positive and FlpoER-negative. **e.** Expression of Cre-dependent (not intersectional) AAV-DIO-mCherry in Htr2a-Cre;Wfs1-FlpoER mice showing the entire population of CeA^Htr2a^ cells in the CeL and CeM. **f.** Expression of AAV-Con/Fon-EYFP in Htr2a-Cre;Wfs1-FlpoER mice showing predominantly the CeL fraction of Htr2a-Cre positive cells. **g,h.** Quantifications of the fraction of Htr2a-Cre::Con/Fon-EYFP positive cells in the CeL (54.3±7.5%) among all CeA^Htr2a^ cells (mCherry+) in CeL **(g)** and of the fraction of Htr2a-Cre;Con/Fon-EYFP positive cells in CeM (20±6.2%) among the total number of CeA cells labelled by the virus **(h)** (n = 3 mice/9 sections). **i.** Expression of Cre-dependent (not intersectional) AAV-DIO-mCherry in Htr2a-Cre;Wfs1-FlpoER mice. **j.** Expression of AAV-Con/Foff-EYFP in Htr2a-Cre;Wfs1-FlpoER mice showing predominantly the CeM fraction of Htr2a-Cre positive cells. **k,l.** Fractions of Htr2a-Cre::Con/Foff-EYFP positive cells in CeM (66.3±2.0%) among all CeA^Htr2a^ cells in CeM **(k)** and of the Htr2a-Cre::Con/Foff-EYFP positive cells in CeL (30.3±4.3%) among all CeA^Htr2a^ cells expressing the virus **(l)** (n = 3 mice/ 9 sections). **m.** Expression of Cre-dependent (not intersectional) AAV-DIO-mCherry in Sst-Cre;Wfs1-FlpoER mice showing the entire population of CeA^Sst^ cells in CeL and CeM. **n.** Expression of AAV-Con/Fon-EYFP in Sst-Cre;Wfs1-FlpoER mice showing predominantly the CeL fraction of Sst positive cells. **o,p.** Quantification of the fraction of Sst-Cre::Con/Fon-EYFP positive cells in CeL (47.5±7.7%) among all Sst positive cells in CeL **(o)** and of the fraction of Sst-Cre::Con/Fon-EYFP positive cells in CeM (8.7±2.3%) among the total number of the cells labelled by the virus **(p)** (n =3 mice/9 sections). **q.** Expression of Cre-dependent (not intersectional) AAV-DIO-mCherry in Sst-Cre;Wfs1-FlpoER mice. **r.** Expression of AAV-Con/Foff-EYFP in Sst-Cre;Wfs1-FlpoER mice showing predominantly the CeM fraction of Sst positive cells. **s,t.** Fractions of Sst-Cre::Con/Foff-EYFP positive cells in CeM (68.6±5%) among all CeA^Sst^ cells in CeM **(s)** and of the Sst-Cre::Con/Foff-EYFP positive cells in CeL (28.7±2.7%) among all CeA^Sst^ cells expressing the virus **(t)** (n = 3 mice/ 9 sections). Values = Mean± SEM. Scale bar: 115 μm.

Crossing Htr2a-Cre or Sst-Cre with Wfs1-FlpoER mice generated double transgenic ‘intersectional mice’ (Htr2a-Cre;Wfs1-FlpoER and Sst-Cre;Wfs1-FlpoER), in which most cells in the CeL should express both Cre and FlpoER, while the vast majority of cells in the CeM should only express Cre and not FlpoER (Fig. 1c). Injection of INTRSECT Boolean vectors^30, 31^ into the CeA of these mice made it possible to specifically manipulate either the CeL (CreON/FlpON virus or Con/Fon in short) or the CeM population (CreON/FlpOFF virus or Con/Foff) (Fig. 1d). To validate the fidelity of this approach, we injected either Con/Fon or Con/Foff EYFP virus into the CeA of Htr2a-Cre;Wfs1-FlpoER mice (together with a Cre-dependent mCherry virus to visualize the entire Cre-positive population). The results with the Con/Fon EYFP virus showed that EYFP+ cells were 4-times as abundant in the CeL than CeM and encompassed the majority (54%) of Htr2a-Cre-positive cells in the CeL (Fig. 1e-h). Conversely, with the Con/Foff virus, EYFP+ cells were 2.3-times as abundant in the CeM than the CeL and expressed in 66% of the Htr2a-Cre-positive cells in the CeM (Fig. 1i-l). Similar results were obtained with Sst-Cre;Wfs1-FlpoER mice. With the Con/Fon EYFP virus, EYFP+ cells were 10-times as abundant in the CeL than CeM and encompassed a large fraction (48%) of the Sst-Cre-positive cells in the CeL (Fig. 1m-p). With the Con/Foff virus, EYFP+ cells were 2.7-times as abundant in the CeM than CeL and encompassed 69% of the Sst-Cre-positive cells in the CeM (Fig. 1q-t). These experiments show that our Cre/Flp intersectional mouse model, combined with INTRSECT viruses, enabled an enrichment of actuator expression in a specific subpopulation within a single CeA subregion.

### Promotion of water intake exclusively by CeM subpopulations

To study the role of different CeA subpopulations in water consumption, we first asked if optogenetic activation of the neurons would be sufficient to promote water uptake. We stereotactically injected the Con/Fon-or Con/Foff-hChR2-EYFP viruses bilaterally into the CeA of Htr2a-Cre;Wfs1-FlpoER mice to express channelrhodopsin (ChR2) in the CeL and CeM subpopulations, respectively (Fig. 2a). Similar INTRSECT viruses expressing only EYFP control protein were used as negative controls. As positive controls, we expressed ChR2 in the entire CeA^Htr2a^ population by including one cohort of Htr2a-Cre mice that was injected with the simple Cre-dependent DIO-hChR2-EYFP virus (Fig. 2a). Similar injections were performed with Sst-Cre;Wfs1-FlpoER and Sst-Cre mice to analyze the CeA^Sst^ subpopulations. The expression of EYFP in different CeA subregions was validated in all animals (Fig. 2b-e). Optic fibers were placed bilaterally over the CeA to photoactivate the cell bodies of the neurons. In a 30-min paradigm, thirsty animals were exposed to water while being bilaterally photoactivated throughout the session. The same experiment was repeated when animals were hydrated. We found that photoactivation of the entire CeA^Htr2a^ and CeA^Sst^ populations was sufficient to promote water consumption by thirsty as well as hydrated mice (Fig. 2f; Extended data Fig. 2a). Surprisingly, among the two subpopulations, it was exclusively the CeM fractions of both cell types that promoted water uptake both in thirsty and hydrated mice (Fig. 2g,h; Extended data Fig. 2b,c).

**Figure 2.**
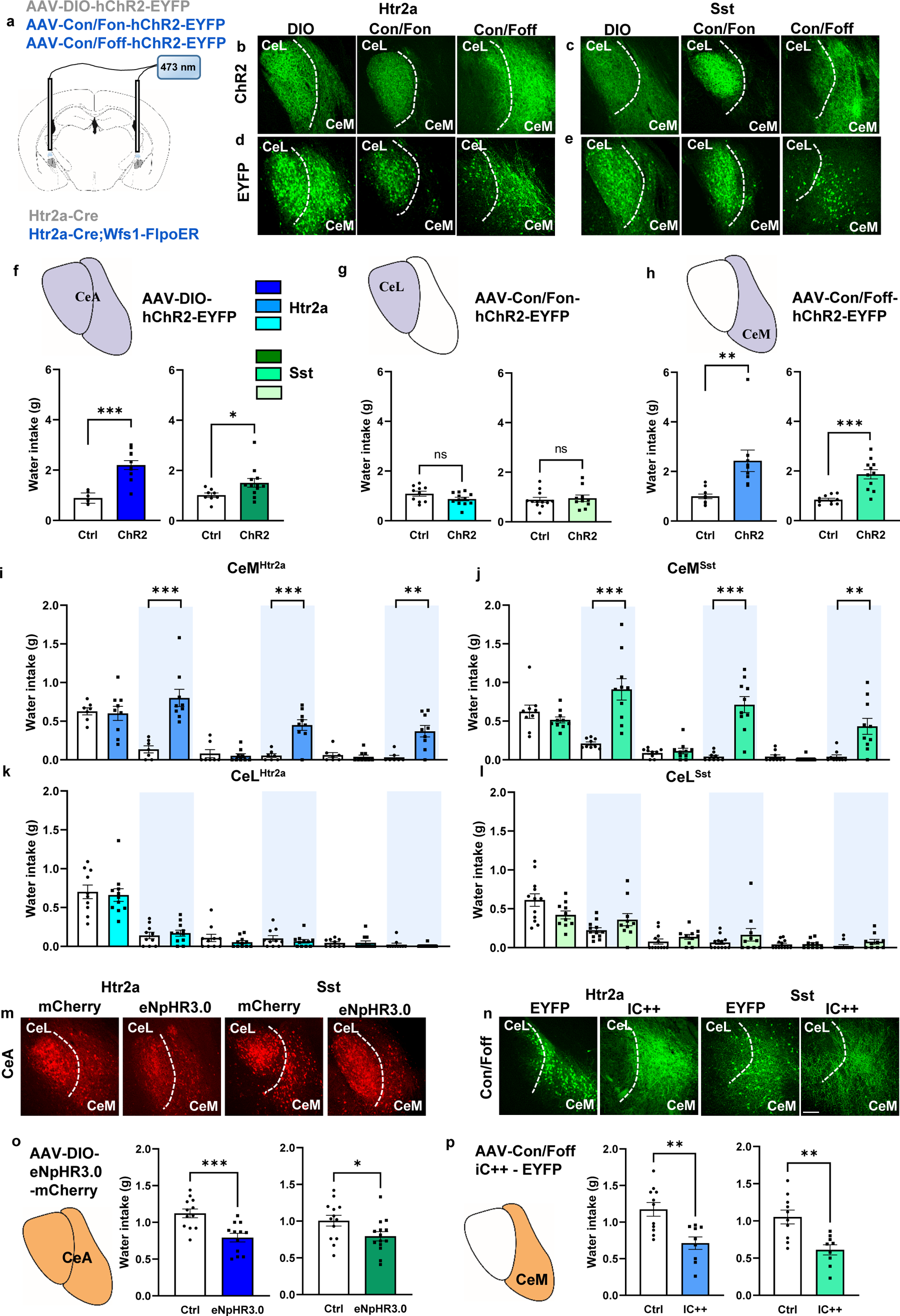
The CeM subpopulations of Htr2a-Cre and Sst-Cre neurons promote water intake. **a.** Scheme representing the viruses injected into the CeA of Htr2a-Cre or Htr2a-Cre;Wfs1-FlpoER mice and optic fiber placement. **b-c.** Representative images of ChR2-EYFP expression in CeA subregions after injections of INTRSECT Con/Fon or Con/Foff viruses into the CeA of Htr2a-Cre;Wfs1-FlpoER **(b**) or Sst-Cre;Wfs1-FlpoER mice **(c)**. **d-e.** Representative images of control EYFP protein expression after similar manipulations described in panels b and c. **f.** Photoactivation of the entire CeA^Htr2a^ or CeA^Sst^ neuron population for 30-minutes in water-deprived mice increased water intake compared to photoactivated control animals (Htr2a: unpaired t test *p*<0.0001, t=6.429. Sst: Mann-Whitney U test *p*=0.0124, U=16). **g.** Photoactivation of the CeL fractions had no effect on water intake (Htr2a: unpaired t test *p*=0.0913, t=1.774. Sst: unpaired t test *p*=0.7017 t=0.3886). **h.** Photoactivation of the CeM fractions of Htr2a-Cre or Sst-Cre mice was sufficient to increase water consumption compared to photoactivated control animals (Htr2a: Mann-Whitney U test *p*<0.0001, U=2. Sst: unpaired t test *p*<0.0001 t=5.054). **i-l.** ChR2-expressing mice were subjected to alternating 10-minutes light OFF and light ON epochs (photoactivation). Photoactivation of CeM^Htr2a^ **(i)** or CeM^Sst^ neurons **(j)** in water deprived mice increased water intake during light ON epochs (Htr2a: 10-20 min Mann-Whitney U test *p*=0.0002, U=0. 30-40 min: unpaired t test *p*=0.0003, t=4.801. 50-60 min: Mann-Whitney U test *p*=0.0030, U=5.500. Sst: 10-20 min unpaired t test *p*=0.0002, t=4.738. 30-40 min: Mann-Whitney U test *p*<0.0001, U=1.500. 50-60 min: Mann-Whitney U test *p*=0.0012, U=8). Photoactivation of the CeL fractions of Htr2a-Cre **(k)** or Sst-Cre neurons **(l)** had no significant effect on water intake (Htr2a: 10-20, 30-40, 50-60 min: unpaired t test or Mann-Whitney U test *p*>0.05. Sst: 10-20, 30-40, 50-60 min: unpaired t test or Mann-Whitney U test *p*>0.05). **m.** Representative images showing expression of Cre-dependent eNpHR3.0 and control mCherry in the entire CeA of Htr2a-Cre and Sst-Cre animals. **n.** Representative images showing expression of IC++ and control YFP predominantly in the CeM after injection of Con/Foff virus in Htr2a-Cre;Wfs1-FlpoER and Sst-Cre;Wfs1-FlpoER mice. **o.** Photoinhibition of the entire populations of CeA^Htr2a^ (blue) or CeA^Sst^ neurons (green) in water deprived mice significantly decreased water consumption compared to photoinhibited controls during a 30-min drinking assay (Htr2a: unpaired t test *p*=0.0007, t=3.939. Sst: unpaired t test *p*=0.0363, t=2.213). **p.** Photoinhibition of the CeM fractions of CeA^Htr2a^ (blue) or CeA^Sst^ neurons (green) also significantly decreased water consumption compared to photoinhibited controls in the same assay (Htr2a: unpaired t test *p*=0.0022, t=3.578. Sst: unpaired t test *p*=0.0017, t=3.733). Values = Mean± SEM. Scale bar: 115 μm.

To ensure consistency in the effects of optogenetic stimulation across multiple trials, we used a different drinking test during which short (10-min) periods of photoactivation alternated with periods of no photoactivation. During an initial 10-min Light-OFF period, water-deprived mice from all experimental groups were found to consume similar amounts of water (Fig. 2i-l). As the experiment progressed, control animals gradually became hydrated and consumed less water, independent of whether the light was ON or OFF. Among the experimental groups, only photoactivation of the entire CeA populations or the CeM subpopulations increased water consumption during each Light-ON period (Fig. 2i,j; Extended Data Fig. 2d,e). Photoactivation of the CeL subpopulations was insufficient to promote water consumption (Fig. 2k,l). Similar results were obtained with hydrated animals (Extended data Fig. 2f-k). Overall consumption of water was highest during Light-ON periods for mice expressing ChR2 in the entire CeA or CeM subpopulations of Htr2a^+^ and Sst^+^ cells, while water consumption was unaffected in photoactivated controls and mice expressing ChR2 in the CeL (Extended Data Fig. 3). In summary, these experiments showed that the CeM subpopulations of Htr2a-Cre and Sst-Cre positive cells were sufficient to drive water consumption.

We next asked if the CeM subpopulations were also required for water consumption. We stereotactically injected AAVs expressing Cre-dependent eNpHR3.0 bilaterally into the CeA of Htr2a-Cre mice to express the inhibitory Halorhodopsin in the entire CeA^Htr2a^ population (Fig. 2m), or similarly INTRSECT Con/Foff IC++, a blue-shifted halorhodopsin ^31, 35^, into Htr2a-Cre;Wfs1-FlpoER mice to express the actuator specifically in the CeM subpopulation (Fig. 2 n). Similar injections were performed with Sst-Cre;Wfs1-FlpoER mice to analyze the CeM^Sst^ subpopulation (Fig. 2m,n). Corresponding Cre-dependent and INTRSECT viruses expressing mCherry or EYFP control proteins were injected as controls (Fig. 2m,n). Animals were water deprived and tested for water consumption for 30 min while being photoinhibited. Inhibition of the entire CeA population, as well as the respective CeM^Htr2a^ and CeM^Sst^ subpopulations significantly decreased water uptake compared to light-stimulated controls (Fig. 2o,p). Together, these experiments reveal that the CeM subpopulations of Htr2a-Cre and Sst-Cre positive neurons are necessary for efficient water consumption in thirsty mice.

### Promotion of food uptake exclusively by CeM^Htr2a^, but not CeM^Sst^, neurons

Next, we focused our attention on feeding behavior. We previously reported that CeA^Htr2a^ neurons stimulated food intake ^5, 36^, but the specific contributions of the CeL and CeM subpopulations had not been explored. Likewise, the roles of the different Sst subtypes in regulating feeding are not completely understood ^7^. We performed a free-feeding assay with *ad libitum* fed animals expressing ChR2 in Htr2a-Cre^+^ or Sst-Cre^+^ neurons in the CeA, CeL, or CeM. As expected, photoactivation of the entire CeA^Htr2a^ neuron population caused a significant increase in food uptake compared to the photoactivated control group (Fig. 3a). Surprisingly, food uptake was exclusively driven by the CeM^Htr2a^, but not the CeL^Htr2a^, subpopulation (Fig. 3b,c). Photoactivation of the entire CeA^Sst^ neuron population or its subpopulations in the CeL and CeM failed to significantly increase food consumption under these conditions (Fig. 3a-c).

**Figure 3.**
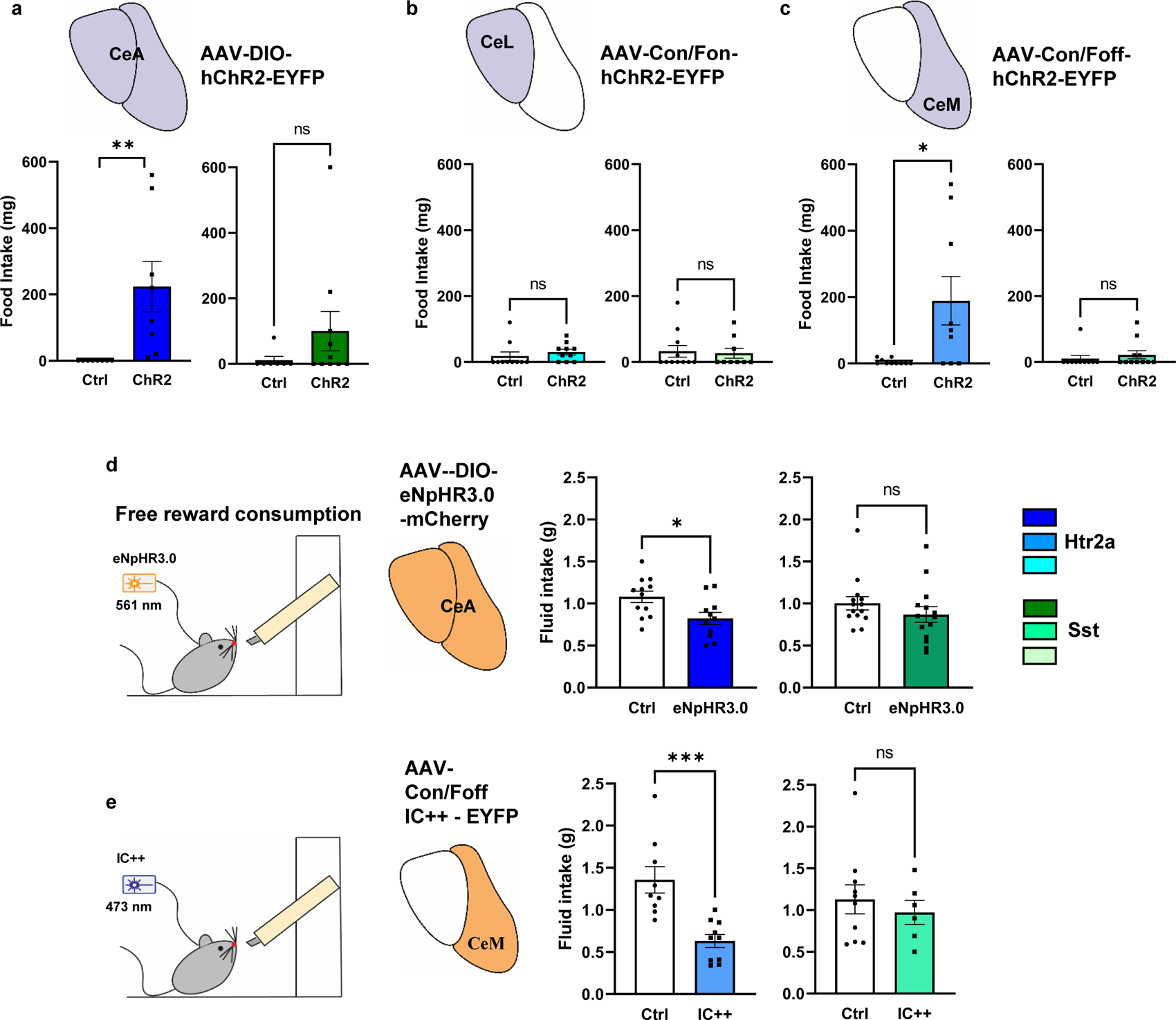
Distinct roles of CeM^Htr2a^ and CeM^Sst^ neurons in feeding behavior. **a.** Photostimulation of the entire population of CeA^Htr2a^ neurons promoted feeding of satiated mice (Wilcoxon signed rank test *p*=0.0078). No statistically significant effect after photostimulation CeA^Sst^ neurons compared to controls (Mann-Whitney U test *p*=0.1821, U=22). **b.** Photostimulation of CeL^Htr2a^ or CeL^Sst^ subpopulations did not promote feeding behavior (CeL^Htr2a^: Mann-Whitney U test *p*=0.1073, U=30.50. CeL^Sst^: *p*=0.9183, U=48). **c.** Photostimulation of CeM^Htr2a^, but not CeM^Sst^ neurons, increased food intake (CeM^Htr2a^: Mann-Whitney U test *p*=0.0235, U=19.50. CeM^Sst^: *p*=0.2783, U=41.50). **d.** Photoinhibition of the entire population of CeA^Htr2a^ neurons in fed-mice decreased the ingestion of a palatable liquid reward compared to controls (unpaired t test *p*=0.0170, t=2.593). No effect by photoinhibition of the entire population of CeA^Sst^ neurons (Mann-Whitney U test *p*=0.1531, U=66.50). **e.** Photoinhibition of CeM^Htr2a^, but not CeM^Sst^, neurons decreased the consumption of a palatable solution compared to controls (CeM^Htr2a^ unpaired t test *p*=0.0005, t=4.313; CeM^Sst^ unpaired t test *p*=0.5433, t=0.6230). Values = Mean± SEM.

To investigate if the activity of the CeM^Htr2a^ subpopulation was required for efficient food consumption, we asked if photoinhibition of CeM^Htr2a^ neurons would reduce consumption of a palatable liquid reward (Fresubin, 2kcal/ml) as previously shown for the entire CeA^Htr2a^ population^5^. Indeed, photoinhibition of the entire CeA^Htr2a^ population or the CeM^Htr2a^ subpopulation significantly reduced the consumption of the palatable liquid compared to the control groups (Fig. 3d,e). In contrast, inhibition of the entire CeA^Sst^ population or the CeM^Sst^ subpopulation did not have a significant effect on Fresubin consumption (Fig. 3d,e). These results provide genetic evidence for CeM^Htr2a^ neurons promoting water and food consumption, whereas CeM^Sst^ neurons had a more restricted role in water, but not food consumption. The CeL subpopulations of Htr2a-Cre^+^ and Sst-Cre^+^ neurons, which highly overlap, do not seem to affect water or food consumption, at least in the assays tested.

### CeM subpopulations drive real-time place preference and conditioned reward behavior

Next, we asked which subpopulations could drive innate rewarding, but non-consummatory behavior, in a real-time place preference (RTPP) assay consisting of a two-chamber arena with one compartment paired with laser photostimulation (Fig. 4a). The results revealed a similar pattern as for water uptake: activation of the entire populations of CeA^Htr2a^ or CeA^Sst^ neurons as well as the respective CeM subpopulations resulted in the animals exhibiting a significant preference for the photostimulation-paired chamber, as compared to the CeL subpopulations and the controls (Fig. 4b-d). The time spent in the center of an open field arena was unchanged, suggesting that the reward behavior in the RTPP assay was not due to an anxiolytic effect (Extended Data Fig. 4a-e).

**Figure 4.**
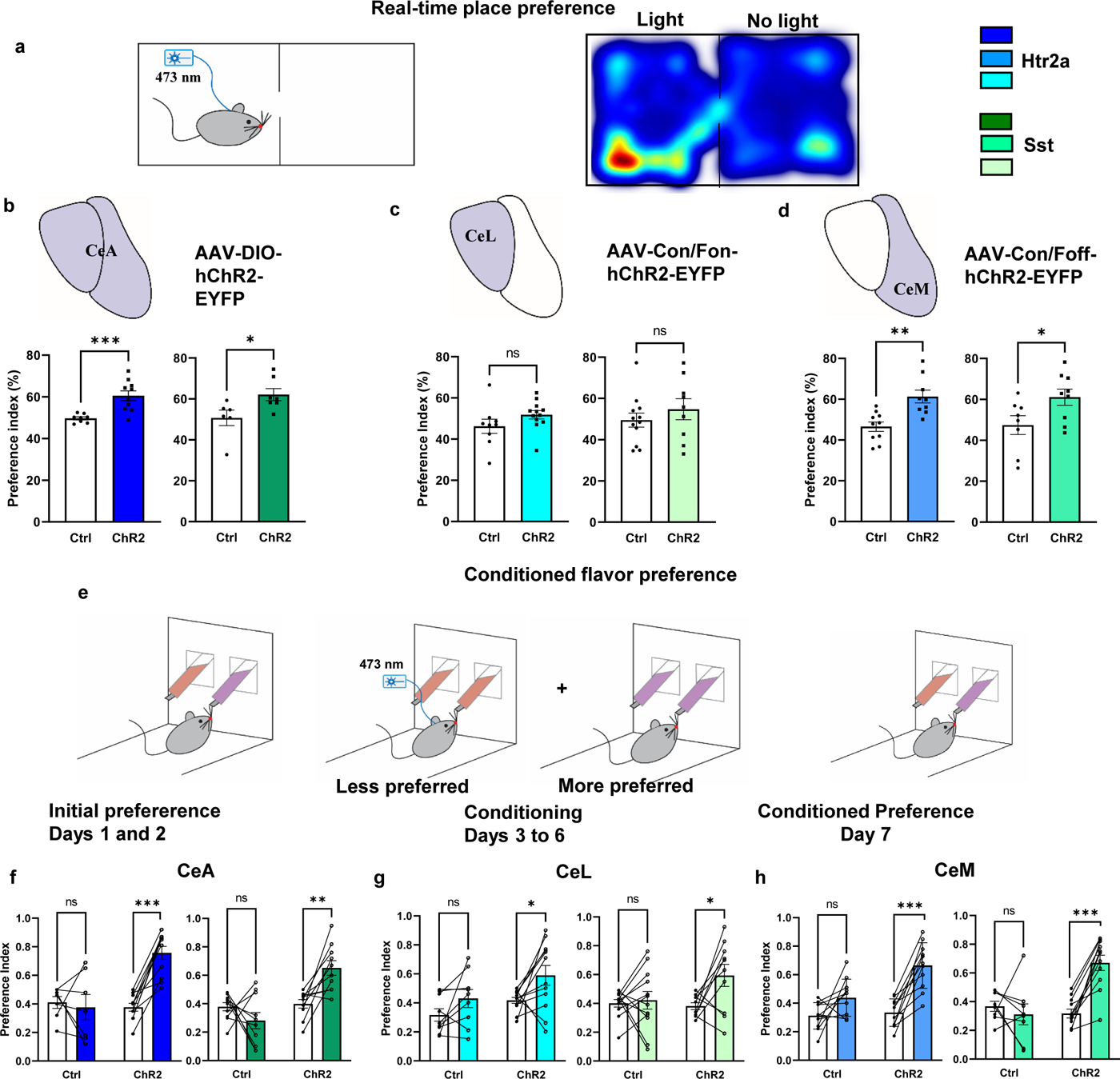
Activation of Htr2a and Sst neurons has a rewarding effect. **a.** Scheme showing the real-time place preference paradigm. Representative heat map of a photostimulated CeM^Htr2a^::ChR2 mouse in the RTPP task. **b.** Preference for the light-paired chamber when the entire populations of CeA^Htr2a^ or CeA^Sst^ neurons are activated in the RTPP task, compared to controls (CeA^Htr2a^: unpaired t test *p*=0.0010, t=4.018. CeA^Sst^: unpaired t test *p*=0.0347, t=2.409). **c.** No preference for the light-paired chamber when only CeL^Htr2a^ or CeL^Sst^ neurons are activated (Htr2a: unpaired t test *p*=0.1521, t=1.492, Sst: unpaired t test *p*=0.3857, t=0.8879). **d.** Preference for the light-paired chamber when only CeM^Htr2a^ or CeM^Sst^ neurons are activated (Htr2a: unpaired t test *p*=0.0013, t=3.844, Sst: unpaired t test *p*=0.0383, t=2.271). **e.** Scheme depicting the conditioned flavor preference test. **f.** Conditioned flavor preference index of mice in which the entire populations of CeA^Htr2a^ (blue) or CeA^Sst^ (green) neurons were photostimulated in comparison to control mice (CeA^Htr2a^: main effect ChR2, Two-way ANOVA, *F*(1,16) =10.12, *p*=0.0058; Bonferroni post-hoc test *p*<0.000. CeA^Sst^: main effect ChR2, Two-way ANOVA, *F*(1,18) =32.06, *p*<0.0001; Bonferroni post-hoc test *p*=0057). **g.** Conditioned flavor preference index of mice in which only the CeL^Htr2a^ (blue) or CeL^Sst^ (green) neurons were photostimulated (CeL^Htr2a^: main effect ChR2, Two-way ANOVA, *F*(1,19) =5.611, *p*=0.0286; Bonferroni post-hoc test *p*=0.0313. CeL^Sst^: main effect Time, Two-way ANOVA, *F*(1,20) =4.918, *p*=0.0383; Bonferroni post-hoc test *p*=0.0255). **h.** Conditioned flavor preference index of mice in which only the CeM^Htr2a^ (blue) or CeM^Sst^ (green) neurons were photostimulated (CeM^Htr2a^: main effect ChR2, Two-way ANOVA, *F*(1,20) =12, *p*=0.0024; Bonferroni post-hoc test *p*<0.0001. CeM^Sst^: main effect ChR2: Two-way ANOVA, *F*(1,17) =8.165, *p*=0.0109; Bonferroni post-hoc test *p*<0.0001). Values = Mean± SEM.

Further, we investigated which subpopulation might condition a preference for a specific flavor. In a reward conditioning paradigm, pairing activation of appetitive CeA neurons with one of two flavors can reverse flavor preference, such that an initially less preferred flavor becomes the preferred one ^5, 17^. Mice expressing ChR2 in the entire populations of CeA^Htr2a^ or CeA^Sst^ neurons, or the respective CeM subpopulations, were allowed to consume two differently flavored non-nutritive liquids. After determining their individual baseline preference, conditioning was performed by pairing the less preferred flavor with optogenetic activation (Fig. 4e). After conditioning, the flavor preference of the mice was assessed by simultaneously offering liquids of both flavors. The results showed that activation of the CeM subpopulations of CeA^Htr2a^ or CeA^Sst^ neurons, reversed the animals’ initial preference, resulting in the least preferred flavor becoming the preferred one (Fig. 4f-h). Somewhat surprisingly, also the CeL subpopulations showed significant conditioning activities (Fig. 4g). Consistent with these results, mice expressing ChR2 in the CeM or CeL subpopulations of CeA^Htr2a^ or CeA^Sst^ neurons consumed a significantly larger amount of the liquid that was paired with light stimulation compared to controls (Extended Data Fig. 4f-h). These results demonstrate that the activities of both CeM^Htr2a^ and CeM^Sst^ subpopulations are intrinsically positively reinforcing in RTPP and conditioned flavor preference assays. Both, CeL^Htr2a^ and CeL^Sst^ subpopulations displayed modest reinforcing activities in the conditioned flavor preference assay, but not in real-time place preference.

### CeM^Htr2a^ and CeM^Sst^ neuron responses to different rewarding behaviours

To understand how CeM^Htr2a^ and CeM^Sst^ neurons participate in consummatory behavior, we performed single-cell resolution *in vivo* calcium imaging in freely behaving mice. We injected a Con/Foff GCaMP6m virus unilaterally into the CeA of either Htr2a-Cre::Wfs1-FlpoER or Sst-Cre::Wfs1-FlpoER animals to record the neuronal activities preferentially from CeM subpopulations. A gradient-index (GRIN) lens was implanted to monitor the neuronal activity using a head-mounted miniscope. We tested the animals in three separate conditions: exposure to water, food, and Fresubin. To make consumption of water and food pleasant, mice were water- and food-deprived, respectively. Fresubin instead is very palatable independent of the hunger state. Hence, mice exposed to Fresubin were fed *ad libitum*. Recordings were done for 10 min following a 10 min habituation period (Fig. 5a,b). During habituation, we recorded the activities of an average of 128 neurons per subpopulation, while during stimulation, the numbers of active neurons increased to an average of 160 neurons (Extended data Fig. 5a). The average number of neurons that were active across different rewarding conditions ranged from 78 to 114 (Extended data Fig. 5b). When comparing the activities of all active cells during habituation and stimulation (including active and inactive times), we observed a general increase in neuronal activities for both CeM^Htr2a^ and CeM^Sst^ neurons during stimulation, except for CeM^Htr2a^ neurons exposed to water (Fig. 5c,d). When comparing the activities during reward consumption versus inactive episodes during the 10-min stimulation periods, we found that CeM^Htr2a^ and CeM^Sst^ neurons showed higher activities during drinking and feeding (Extended data Fig. 5c).

**Figure 5.**
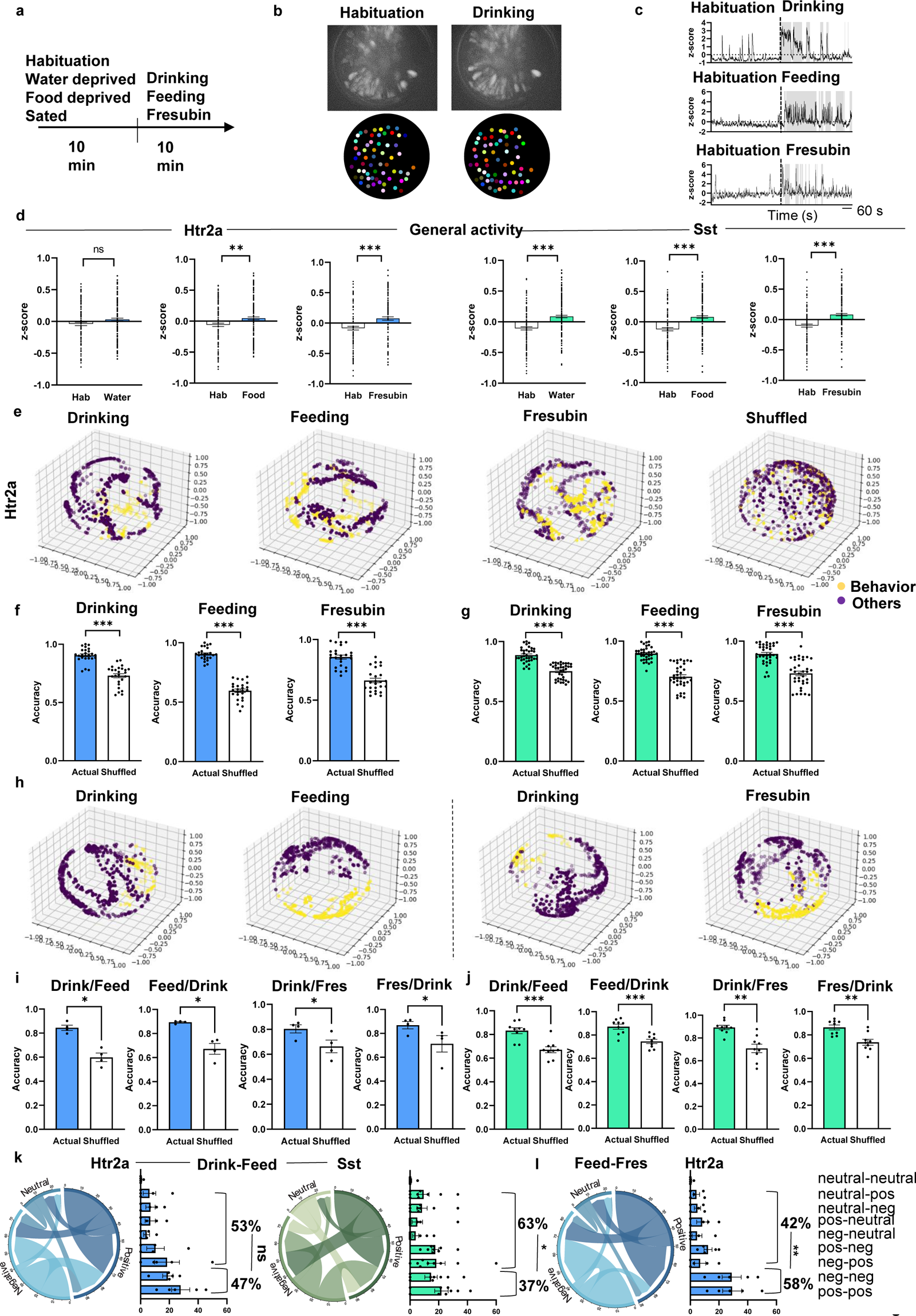
CeM^Htr2a^ and CeM^Sst^ neurons are differentially activated by multiple rewarding stimuli. **a.** Scheme depicting the behavioral paradigm. **b.** Maximum projection images during 10-minute habituation and 10-minute exposure to water of a representative Htr2a-Cre;Wfs1-FlpoER mouse injected into the CeM with Con/Foff-GCaMP6m to record calcium responses from CeM^Htr2a^ neurons. **c.** Representative traces from CeM^Htr2a^ neurons showing increased neuronal activities during drinking, feeding and Fresubin consumption. Grey color indicates consumption bouts. **d.** Average z-score comparisons between periods of inactivity and bouts of reward consumption of CeM^Htr2a^ and CeM^Sst^ neurons (Wilcoxon matched-pairs signed rank test. CeM^Htr2a^: drinking, *p*=0.0079; feeding, *p*<0.0001; Fresubin, *p*=0.4580. CeM^Sst^: drinking, *p*=0.0008; feeding, *p*=0.0052; Fresubin, *p*=0.0065). **e.** 3D Visualization of CEBRA generated embeddings of CeM^Htr2a^ neurons considering consumption behavior and neural dynamics over time. Shuffled: CEBRA embeddings by CeM^Htr2a^ drinking shuffled data. The three dimensions represent the three CEBRA principle components. Each dot represents a time point. Yellow dots mark the time the animals are consuming rewards; purple dots the time the animal is not involved in consumption. **f,g.** Comparison of behavioral decoding accuracy using a Random Forest (RF) algorithm across animals for CeM^Htr2a^ neurons **(f)** (drinking, feeding, Fresubin: : Wilcoxon matched-pairs signed rank test *p*<0.0001) and CeM^Sst^ neurons **(g)** (drinking, feeding, Fresubin: : Wilcoxon matched-pairs signed rank test *p*<0.0001). **h.** CEBRA generated embeddings of a subset of CeM^Htr2a^ neurons that were longitudinally detected during drinking and feeding sessions and during drinking and Fresubin consumption. **i,j.** Accuracy of behavioral decoding through RF, employing CEBRA embeddings of CeM^Htr2a^ **(i)** and CeM^Sst^ **(j)** neurons longitudinally detected during drinking and feeding or drinking and Fresubin consumption. The embeddings derived from drinking behavior were used for predicting feeding and those from feeding for predicting drinking. Furthermore, the drinking embeddings were applied to predict Fresubin consumption and vice versa (paired t test. CeM^Htr2a^: drinking-feeding: *p*=0.0158, t=4.956; feeding-drinking: *p*=0.0175, t=4.768; drinking-Fresubin: *p*=0.0292, t=3.935; Fresubin-drinking: *p*=0.0337, t=3.724. CeM^Sst^: drinking-feeding: *p*<0.0001, t=6.605; feeding-drinking: *p*=0.0002, t=6.044; drinking-Fresubin: *p*=0.0010, t=4.763; Fresubin-drinking: *p*=0.0009, t=4.850). **k.** Chord diagrams and bar graphs depicting the stable or unstable correlation of the same CeM^Htr2a^ and CeM^Sst^ neurons detected during drinking and feeding sessions. (CeM^Htr2a^ stable v.s. unstable: unpaired t test *p*=0.5418, t=0.6371. CeM^Sst^ stable v.s. unstable: unpaired t test *p*=0.0193, t=2.602). **l.** Chord diagram and bar graph depicting the stable or unstable correlation of the same CeM^Htr2a^ neurons detected during feeding and Fresubin sessions. (CeM^Htr2a^ stable v.s. unstable: unpaired t test *p*=0.0017, t=4.627). Values = Mean± SEM.

To investigate whether consummatory behaviors contributed to the dynamics of neural activity, we analyzed the data using the recently developed machine learning algorithm CEBRA^37^ (for further details, see Methods). We applied the so-called Hybrid model, which considers the relation between consumption bouts and the patterns of neural activities across time. If a behaviour contributed significantly to neural activity, we expected to see clear structure in CEBRA-generated 3D neural embeddings compared to shuffled embeddings. When applying CEBRA to both CeM^Htr2a^ and CeM^Sst^ calcium recordings during drinking, feeding and Fresubin consumption, we found clear structures for all three behaviors, i.e. a separation of data points derived from times when the animals were engaged in consummatory behavior from times when the animals were not involved in consumption (Fig. 5e). Notably, this separation was lost when the data was shuffled, suggesting contributions of these behaviours to neural activities (Fig. 5e and data not shown). To analyze how well the neural embeddings decoded specific behavioral features, we applied a Random Forest (RF) classifier for behavioral decoding (see Methods). The results indicated that the CeM^Htr2a^ and CeM^Sst^ neural embeddings decoded the respective behaviours with 85-90% accuracy, significantly better than shuffled data (60-75%) (Fig. 5f,g). To measure the prediction error of the RF classifier, we calculated the out of bag (OOB) error, and found that in all cases the OOB error from the actual data was much lower than from shuffled data (Extended data Fig. 5d). Next, we asked if neurons associated with different behaviors were preserved across contexts with shared activities, using longitudinal registration of recorded neurons. We applied CEBRA to neural activity data of neurons shared between two behaviours (e.g. drinking/feeding, drinking/Fresubin). We found that the neurons associated with both behaviours showed clearly structured CEBRA embeddings (Fig. 5h, Extended data Fig. 5e). The neural embeddings decoded the behavioral predictions with higher accuracy compared to the results obtained with shuffled data (Fig. 5i,j, Extended data Fig. 5f).

Through correlation analysis between behavior and neuronal activity, we found ensembles of CeM^Htr2a^ and CeM^Sst^ populations whose activities correlated positively and negatively with reward consumption (Extended Data Fig. 5g). The CeM^Sst^ populations responded rather homogeneously to stimulation, with 46-48% of cells displaying positive correlation with all three rewards (Extended Data Fig. 5h). The responses of CeM^Htr2a^ neurons were more heterogenous, with 39-59% showing positive correlation (Extended Data Fig. 5h). The highest positive correlation of CeM^Htr2a^ neurons was with feeding (59%), consistent with their observed function in food consumption.

Next, we asked how the activities of individual cells changed between the rewards having different physical properties. We compared all cells that were active during two behavioral sessions, e.g. drinking and feeding or feeding and Fresubin, and asked which fractions of cells kept their correlated activities constant (positive – positive, or negative – negative correlation) versus fractions that changed their activities (Fig. 5k,l; Extended data Fig. 5 j). When comparing drinking and consuming solid food, the fraction of CeM^Htr2a^ cells displaying a stable correlation (pos – pos, neg – neg) was comparable to the fraction of cells that changed their activities (47% versus 53%) (Fig. 5k). Instead, the fraction of CeM^Sst^ cells that changed their activities was significantly larger than the fraction of cells displaying a stable correlation (63% versus 37%) (Fig. 5k). When comparing solid food and liquid Fresubin consumption, the fraction of CeM^Htr2a^ cells displaying a stable correlation was significantly larger than the fraction of cells that changed their activities (58% versus 42%) (Fig. 5l). Instead, the fraction of CeM^Sst^ cells showing a stable correlation was comparable to the fractions of cells that changed their activities (Extended Fig. 5j). In summary, at the population level, the majority of CeM^Htr2a^ and CeM^Sst^ neurons increased their activities during reward consumption and their activity dynamics contributed to and decoded specific behavioral features. Depending on the physical attributes of the rewards, CeM^Htr2a^ and CeM^Sst^ neurons were recruited into different ensembles with constant or variable correlated activities.

### CeM^Htr2a^ and CeM^Sst^ neurons respond differently to stimuli of opposite valence

To investigate what fractions of CeM^Htr2a^ and CeM^Sst^ neurons respond specifically to one class of stimuli (“specializers”) or exhibit a general response to a broad range of stimuli even of opposite valence (“generalizers”), we exposed the same cohorts of mice expressing GCaMP6m to three different aqueous solutions: water, saccharin and the bitter tastant quinine. During each 10-minute session, the stimulus was orally administered at minutes 1 and 6 by an experimenter who the mice were well accustomed to. During the remaining time, mice were allowed to freely behave (Fig. 6a). We recorded from an average of 107 CeM^Htr2a^ and 101 CeM^Sst^ neurons and an average of 71 neurons was longitudinally detected in different sessions (Extended data Fig. 6a,b). Many CeM^Htr2a^ and CeM^Sst^ neurons showed an increase in calcium responses to all three stimuli during the stimulation periods (Fig. 6b). When analyzing the average population activities across multiple 10-min sessions, we found that the switch from water to saccharin, and water to quinine-added water, did not significantly change activities (although the latter was close to significance). Interestingly, we observed a significant decrease in population activity when switching from saccharin to quinine (Fig. 6c). These results suggest that the switch from sweet to bitter taste quenched the activities of both subpopulations. We applied the CEBRA Hybrid model on the recorded neuronal activities of both CeM^Htr2a^ and CeM^Sst^ neurons during the water, saccharin, and quinine behaviors. The observable structures suggested that these behaviors contributed significantly to the alterations in neuronal activities (Fig. 6d). The embeddings of both neuron populations could accurately decode positive and negative stimuli with higher accuracies (97-100%) and lower OOB errors compared to the shuffled data (71-73%) (Fig. 6e; Extended Data Fig. 6d).

**Figure 6.**
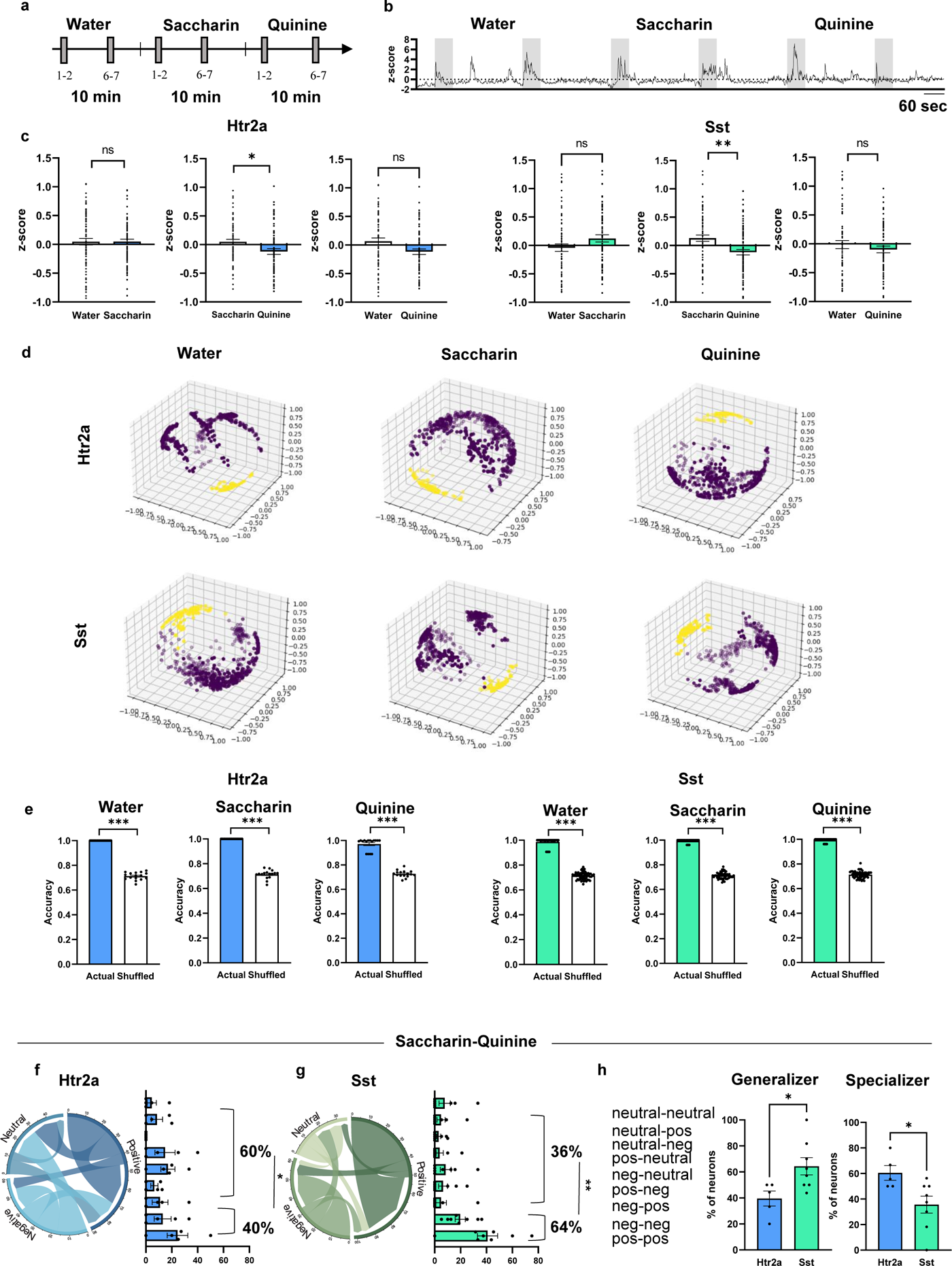
Different salient stimuli elicit responses in CeM^Htr2a^ and CeM^Sst^ neurons. **a.** Scheme describing the behavioral paradigm. Water-deprived mice were exposed to three 10-min long consecutive sessions in which they were administered with water, saccharin, or quinine at minutes one and six of each 10-min interval. **b.** Representative trace from a CeM^Htr2a^ neuron showing increased neuronal activity during most of the 1min consumption bouts (grey color). **c.** Average z-score comparisons of the activities of the same CeM^Htr2a^ and CeM^Sst^ neurons recorded in the three different conditions (Wilcoxon matched-pairs signed rank test. CeM^Htr2a^: water-saccharin, *p*=0.9477; saccharin-quinine, *p*=0.0175; water-quinine, *p*=0.0714. CeM^Sst^: water-saccharin, *p*=0.1836; water-quinine, *p*=0.6808; saccharin-quinine, paired t test *p*=0.0067, t=2.788). **d.** CEBRA generated embeddings of CeM^Htr2a^ and CeM^Sst^ neurons under water, saccharin and quinine conditions. **e.** Comparison of behavioral decoding accuracy using Random Forest across animals for Htr2a (in blue) (water,saccharin,quinine: Wilcoxon matched-pairs signed rank test *p*<0.0001) and Sst (green) (water,saccharin,quinine: : Wilcoxon matched-pairs signed rank test *p*<0.0001) neurons. **f,g.** Chord diagrams and bar graphs showing the stable or unstable correlation of the same CeM^Htr2a^ **(f)** (unpaired t test *p*=0.0332, t=2.569) and CeM^Sst^ **(g)** (unpaired t test *p*=0.0087, t=3.047) neurons from saccharin to quinine. **h.** Percentages of generalizers and specializers within the CeM^Htr2a^ and CeM^Sst^ populations in the saccharin-quinine comparison (Generalizer: unpaired t test *p*=0.0259, t=2.573. Specializer: unpaired t test *p*=0.0261, t=2.570). Values = Mean± SEM.

Next, we compared the activities during the stimulation episodes with the times in-between and asked which cells were positively or negatively correlated, or showed an activity that was not significantly correlated with stimulation (neutral). We then asked how many cells could be qualified as generalizers showing a stable correlation (positive-positive, negative-negative) between oppositely valenced stimuli, and how many cells would be specializers that switched their correlation. For CeM^Htr2a^ neurons, during the switch from saccharin to quinine, the fraction of specializers was significantly larger than the fraction of generalizers (60% versus 40%). Conversely, for CeM^Sst^ cells the pattern was opposite, with generalizers outnumbering specializers (64% versus 36%) (Fig. 6 f-h,). Other comparisons did not yield significant differences between generalizers and specializers (Extended Fig. 6 e). In summary, these results suggest that the activities of CeM^Htr2a^ and CeM^Sst^ neurons contribute to the detection of stimuli of opposite valence, and that the CeM^Htr2a^ population contains more cells that specialize in encoding valence-specific stimuli than CeM^Sst^ neurons.

### Appetitive CeM neurons are inhibited by anorexigenic CeA^PKCδ^ neurons

Next, we asked how the activities of appetitive CeM^Htr2a^ and CeM^Sst^ may be regulated. Previous studies had suggested that putative appetitive CeA neurons may be under inhibitory control of anorexigenic CeA^PKCδ^ neurons ^6^ and that CeA^Htr2a^ neurons engage in reciprocal inhibitory connections with CeA^PKCδ^ neurons ^5^. This raised the possibility that CeM^Htr2a^ and CeM^Sst^ neurons may also receive direct inhibitory input from CeA^PKCδ^ neurons that reside in the CeL/C subregion. We hypothesized that inhibition of CeA^PKCδ^ neurons would disinhibit and thereby activate appetitive CeM^Htr2a^ and CeM^Sst^ neurons, whereas activation of CeA^PKCδ^ neurons would inhibit them. We first confirmed that photoactivation of PKCδ cells suppressed water consumption compared to photostimulated control mice (Fig. 7 a-d). In a 10-minute ON/OFF stimulation protocol (similar to Fig. 2, but starting with Light ON), the control group consumed most of the water during the initial 10 minutes, then gradually decreased consumption as the mice became satiated, regardless of the light phase. In contrast, CeA^PKCδ^ neurons consumed water only during the Light OFF phases (Fig. 7 d). Conversely, photoinhibition of CeA^PKCδ^ neurons using Cre-dependent Halorhodopsin (Fig. 7 e,f) resulted in increased water uptake compared to photostimulated control mice (Fig. 7 g). Notably, anxiety did not appear to affect the behavioral outcomes, as the mice subjected to photo-activation or inhibition of CeA^PKCδ^ neurons spent a comparable amount of time in the center-zone during the open field test as the control group (Fig. 7 h,i).

**Figure 7.**
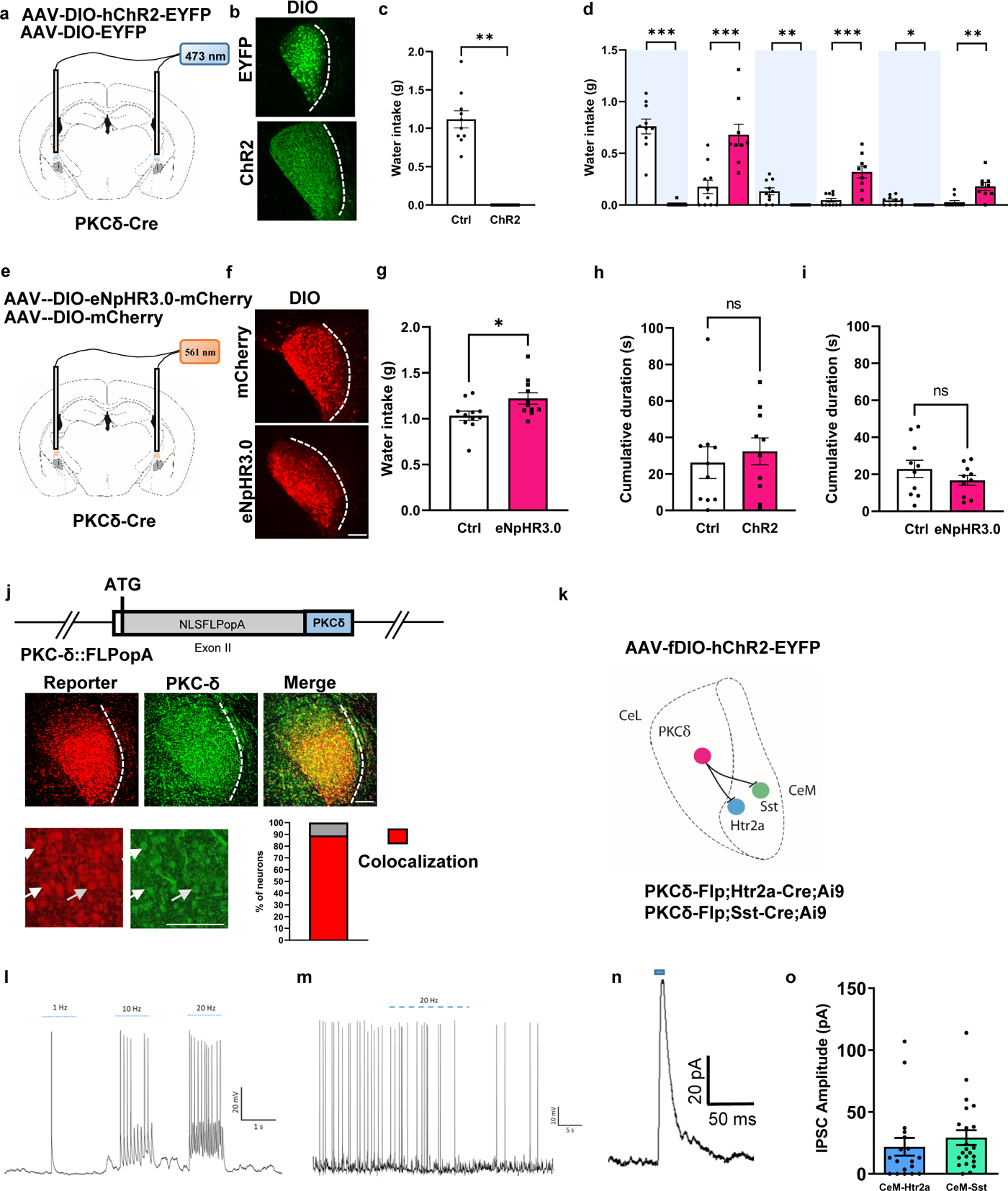
Inhibition of appetitive CeM neurons by thirst quenching CeA^PKCδ^ neurons. **a.** Scheme representing the viruses injected bilaterally into the CeA of PKCδ-Cre mice and optic fiber placement. **b.** Representative images of EYFP and ChR2 expression in the CeL of PKCδ-Cre mice. **c.** Activation of CeA^PKCδ^ neurons for 30 min completely abolished water uptake by water-deprived mice (Wilcoxon signed rank test *p*=0.0020). **d.** Water-deprived mice expressing ChR2 or EYFP in CeA^PKCδ^ neurons, were exposed to water in a 10 min Light ON/OFF behavioral paradigm (Light ON is indicated by light blue shading). While photoactivation of CeA^PKCδ^ cells drastically diminished drinking behavior during the first 10-min light ON epoch, control mice were not affected by the light stimulation and consumed the majority of the water during this time period. In contrast, PKCδ-Cre;ChR2 mice consumed water only during Light OFF epochs (0-10 min Mann-Whitney U test *p*<0.0001, U=0; 10-20 min Mann-Whitney U test *p*=0.0004, U=5; 20-30 min Wilcoxon signed rank test *p*=0.0078; 30-40 min Mann-Whitney U test *p*=0.0002, U=4; 40-50 min Wilcoxon signed rank test *p*=0.0313; 50-60 min Mann-Whitney U test *p*=0.0011, U=9). **e.** Scheme depicting the eNpHR3.0-mCherry and control mCherry viruses injected bilaterally into the CeA of PKCδ -Cre mice and optic fiber placement. **f.** Representative images of mCherry and eNpHR3.0-mCherry expression in the CeL of PKCδ-Cremice. **g.** Photoinhibition of PKCδ neurons significantly increased water uptake in water-deprived mice (unpaired t test *p*=0.0282, t=2.365). **h,i.** Time spent in the center zone during an open-field test by photoactivated PKCδ::ChR2 and PKCδ::EYFP mice (**h**) (Mann-Whitney U test *p*=0.4359, U=39) and by photoinhibited PKCδ::eNpHR3.0 and PKCδ::mCherry mice (**i**) (unpaired t test *p*=0.2719, t=1.133). **j.** Scheme illustrating the new PKCδ-Flp transgene. NLSFLPopA, optimized FLP carrying a nuclear localization sequence and mRNA transcript with polyA recognition sequence. Validation of Flp expression. Flp-mediated recombination in PKCδ-Flp;FPDi mice leads to expression of mCherry (red) in the CeA. In green endogenous PKCδ immunoreactivity. Fraction of mCherry+ among PKCδ immunopositive cells was 89%; n=3 brains, more than 3 sections per brain. **k.** Scheme representing the intersectional strategy to demonstrate the functional connection between PKCδ and CeM^Htr2a^ cells using PKCδ-Flp;Htr2a-Cre;Ai9 mice. CeA^PKCδ^ neurons express a Flp-dependent ChR2-EYFP virus and CeM^Htr2a^ cells are TdTomato positive. Similar strategy for the connection between PKCδ and CeM^Sst^ cells. **l.** Representative example of a CeA^PKCδ^ neuron expressing ChR2 whose firing is induced by blue light stimulation (1 Hz, 10 Hz, 20 Hz) in slice recordings. **m.** Representative example of a photostimulated (20 Hz) CeA^PKCδ^ neuron suppressing current-injection-induced firing of a CeM^Sst^ neuron. **n.** Representative trace of induced inhibitory postsynaptic current (IPSC) showing the response in CeM^Sst^ cells after photoactivation of CeA^PKCδ^ neurons. **o.** Quantification of the IPSC amplitudes of the recorded CeM^Htr2a^ and CeM^Sst^ neurons (n=22 for CeM^Htr2a^, n=18 for CeM^Sst^). Values = Mean± SEM. Scale bar: 115 μm.

To demonstrate a functional connection between CeA^PKCδ^ and the appetitive CeM subpopulations, we generated a new mouse line PKCδ-Flp, that expresses the Flp recombinase specifically under the control of the PKCδ promoter (Fig. 7 j). We validated the expression of the Flp recombinase by crossing the PKCδ-Flp mouse line with a Flp-dependent reporter mouse and quantified the numbers of reporter-positive cells versus PKCδ immunostaining. The results revealed that 89% of the PKCδ immunopositive cells colocalized with the reporter (Fig. 7 j). Using an intersectional approach, we crossed PKCδ-Flp mice with Htr2a-Cre mice also carrying a Cre-dependent tdTomato reporter (Ai9) to later visualize CeM^Htr2a^ neurons in slices. Similar crosses were done with Sst-Cre mice (Fig. 7 k). We then injected a Flp-dependent ChR2-EYFP AAV into the CeA to photoactivate CeA^PKCδ^ neurons in slices. With this design, it was possible to patch and record from CeM tdTomato-positive neurons while photoactivating CeA^PKCδ^ neurons in the CeL (Fig. 7 k). These results show that photoactivation of CeA^PKCδ^ neurons suppresses current-induced firing of both Htr2a and Sst cells in the CeM (Fig. 7 l,m and data not shown). Additionally, we could record inhibitory post-synaptic currents from CeM^Htr2a^ and CeM^Sst^ neurons, providing evidence for a monosynaptic connection from CeA^PKCδ^ to both CeM subpopulations (Fig. 7 n,o). Together, these findings demonstrate that CeA^PKCδ^ neurons suppress water uptake and inhibit the activities of CeM^Htr2a^ and CeM^Sst^ neurons in slices, consistent with a model in which the activities of CeM^Htr2a^ and CeM^Sst^ neurons are under inhibitory control of CeA^PKCδ^ neurons *in vivo*.

### Htr2a and Sst neurons send projections to brain regions associated with reward processing

After having identified one of the possible mechanisms regulating the activities of Htr2a and Sst neurons in the CeM, we proceeded to anatomically map the long-range outputs of these neurons. To identify the major output targets of Htr2a and Sst neurons in the CeL and CeM, we selectively expressed either Con/Fon or Con/Foff YFP viruses in Htr2a/Sst-Cre::Wfs1-FlpoER mice. As expected, our analysis revealed that the CeM populations show a larger number of outputs compared to the CeL populations (Fig. 8). Moreover, in the regions where CeL and CeM populations shared common outputs, projections from the CeM often covered a larger area within the target region. This included projections to the bed nucleus of the stria terminalis (BNST), the interstitial nucleus of the posterior limb of the anterior commissure (IPAC), the parabrachial nucleus (PBN), the substantia nigra (SN), and the periaqueductal gray (PAG) (Fig. 8 a-c; Extended data Fig. 7 a-f). In the PBN, the projections of both CeM^Htr2a^ and CeM^Sst^ neurons covered a broad portion of the PBN including both the lateral and the medial region. In contrast, the projections of both CeL subpopulations were more confined to the lateral region (Fig. 8 a,b). The majority of the brain regions targeted by Htr2a and Sst neurons were similar, while few brain regions were only targeted by one of the cell populations.

**Figure 8.**
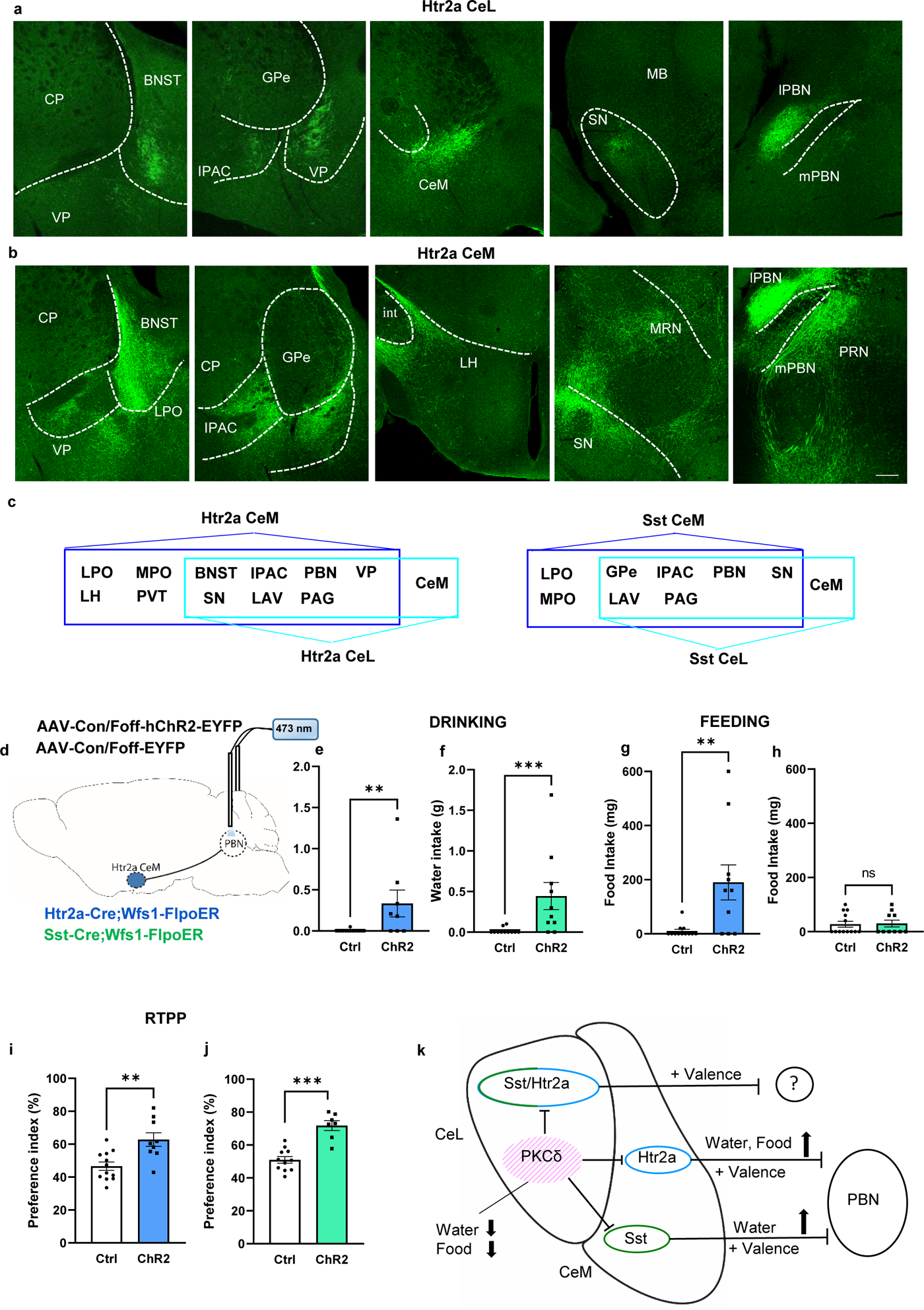
Output regions of CeM^Htr2a^ neurons and functional analysis of CeM→PBN projectors. **a,b.** Representative images showing the brain regions that receive projections from CeL^Htr2a^ (**a**) and CeM^Htr2a^ neurons (**b**). Htr2a-Cre;Wfs1-FlpoER mice were injected either with a Con/Fon-EYFP to target CeL^Htr2a^ or with a Con/Foff-EYFP virus to target CeM^Htr2a^ neurons. Scale bar: 139 μm. **c.** Summary diagrams of the various projections from Htr2a and Sst subpopulations. (BNST: bed nucleus of the stria terminalis. CeM: central amygdala medial part. GPe: globus pallidus. IPAC: interstitial nucleus of the anterior commissure. LAV: lateral vestibular nucleus. LH: lateral hypothalamus. LPO: lateral preoptic area. MPO: medial preoptic area. PAG: periaqueductal gray. L/mPBN: lateral/medial parabrachial nucleus. PVT: paraventricular nucleus of the thalamus. SN: substantia nigra. VP: ventral pallidum). **d.** Scheme depicting the Con/Foff ChR2-EYFP and control viruses injected bilaterally (not shown) into the CeA of Htr2a-Cre;Wfs1-FlpoER or Sst-Cre;Wfs1-FlpoER mice and bilateral (not shown) optic fiber placement above the PBN. **e,f.** Photoactivation of the PBN projections of CeM^Htr2a^ **(e)** or CeM^Sst^ **(f)** significantly stimulated water uptake in normally hydrated mice compared to controls (CeM^Htr2a^: Mann-Whitney U test *p*=0.0021, U=22.50 and CeM^Sst^: *p*=0.0008, U=13). **g,h.** Photoactivation of the PBN projections of CeM^Htr2a^ **(g)**, but not CeM^Sst^ **(h)** significantly stimulated food intake in satiated mice compared to controls (CeM^Htr2a^: Mann-Whitney U test *p*=0.0061, U=23 and CeM^Sst^: *p*=0.9265, U=63.50). **i,j.** Preference for the light-paired chamber when the PBN projections of CeM^Htr2a^ **(i)** or CeM^Sst^ **(j)** are activated in the RTPP task, compared to controls (CeM^Htr2a^ : unpaired t test *p*=0.0023, t=3.522 and CeM^Sst^ : *p*<0.0001 t=5.949). Values = Mean± SEM. **k.** Schematic model of the CeA appetitive microcircuit. The CeL subpopulations of Htr2a and Sst neurons are largely overlapping, and do not promote water or food intake. The CeM subpopulations of Htr2a and Sst neurons are largely distinct (Peters et al., 2023). Both CeM subpopulations promote water intake, while only CeM^Htr2a^ neurons promote feeding. All CeL and CeM appetitive neurons encode positive valence and form long-range projections. The CeM subpopulations of Htr2a and Sst neurons are under inhibitory control of anorexigenic CeA^PKCδ^ neurons which suppress food and water intake. Activation of PBN projections of CeM^Htr2a^ or CeM^Sst^ neurons is sufficient to promote food and water intake, possibly by inhibition of an anorexigenic microcircuit in the PBN.

Many of these regions are known to be involved in rewarding and consummatory behaviors. The BNST plays a central role in reward-related behaviors ^38^, while also influencing feeding through its projections to the lateral hypothalamus (LH) ^39,22,23^. The IPAC, located within the extended amygdala, has been shown to regulate energy homeostasis ^40^. Activation of CeA projections to the SN has been associated with positive reinforcement and the facilitation of learned behaviors in response to salient stimuli ^41, 42^. CeA projections to the PBN modulate food ^5^ and water intake ^43^, as well as alcohol consumption ^23, 44^.

### The appetitive activities of CeM^Htr2a^ and CeM^Sst^ neurons are mediated by projections to the PBN

Based on the known functions of the PBN serving as a hub for sensory information relevant to food and water intake, receiving inputs from interoceptive and exteroceptive sources ^45–51^, and driving aversive emotional behaviors, we hypothesized that inhibition of PBN neurons by CeM^Htr2a^ or CeM^Sst^ neurons may promote drinking and/or feeding behavior. To explore the functions of these neuronal projections, we injected a Con/Foff ChR2 virus, or similar control virus, bilaterally into the CeA of Htr2a-Cre::Wfs1-FlpoER or Sst-Cre::Wfs1-FlpoER mice and placed optic fibers bilaterally above the PBN (Fig. 8d, Extended Data Fig. 8a). Photoactivation of the presynaptic terminals in the PBN of CeM^Htr2a^ or CeM^Sst^ neurons led to increased drinking in both water-deprived and hydrated mice during both a 30 min and a 10 min Light ON/OFF stimulation protocol (Fig. 8e,f, Extended Data Fig. 8c-h.). Furthermore, photoactivation of the CeM^Htr2a^→PBN projectors stimulated feeding behavior, whereas, as expected, no effect was observed for the CeM^Sst^→PBN projectors (Fig. 8g,h). Moreover, both CeM^Htr2a^→PBN and CeM^Sst^→PBN projectors promoted rewarding behavior in real-time place preference (Fig. 8i,j) and conditioned flavor preference assays (Extended Data Fig. 8i-l). No changes in locomotion or anxiety were observed in the open-field test (Extended Data Fig. 18m,n). These findings suggest that CeM^Htr2a^ and CeM^Sst^ neurons promote appetitive and reward behavior through inhibition of PBN neurons.

## DISCUSSION

In this report, we have used an intersectional genetics approach to independently target four putative appetitive neuron subpopulations in the central amygdala: two in the CeL and two in the CeM. Unexpectedly, considering previous reports, we found that neurons mediating water and/or food consumption are confined to the CeM. Separate CeM subpopulations exist for water only (CeM^Sst^), and water or food consumption (CeM^Htr2a^). All four subpopulations are intrinsically positively reinforcing in a conditioned flavor preference assay, but only the two CeM subpopulations mediate real-time place preference. Through *in vivo* calcium imaging we observed that the majority of CeM^Htr2a^ and CeM^Sst^ neurons increased their activities during reward consumption and that their activity dynamics contributed to and decoded specific behavioral features. Depending on the type of rewards, CeM^Htr2a^ and CeM^Sst^ neurons were recruited into different ensembles with constant or variable correlated activities. Calcium imaging further suggests that the activities of CeM^Htr2a^ and CeM^Sst^ neurons contribute to the detection of stimuli of opposite valence, and that the CeM^Htr2a^ population contains more cells that specialize in encoding valence-specific stimuli than CeM^Sst^ neurons. At the microcircuit level, the activities of appetitive CeM neurons are controlled by inhibitory signals from CeA^PKCδ^ neurons and, in turn, appetitive CeM neurons form long-range inhibitory projections to the PBN to promote appetitive and reward behavior. In summary, this study provides a comprehensive functional characterization of molecularly- and anatomically-defined CeA neurons and their roles in appetitive behaviors.

Intersectional genetics using two recombinases has become a valuable tool to address specific neuron subtypes within a larger population of neurons, for example, distinct subtypes of midbrain dopaminergic neurons or hypothalamic POMC neurons ^52, 53^. This method has also been useful to manipulate specific neurons within an anatomically well-defined area, for example, specific Cre+ neuron populations in the spinal cord, and to assess the functional consequences without interference by Cre recombination in the brain ^54^. Here, we have used a newly generated mouse line that expresses the optimized and tamoxifen-inducible FlpoER recombinase in the CeL subdivision of the CeA, to a much higher degree (17-fold) than in cells of the CeM. In combination with a specific Cre line and a Con/Fon Boolean reporter virus, we could target between 4- and 10-times more Cre+ cells in the CeL than CeM. Conversely, with a Con/Foff virus, we could target between 2.3- and 2.7-times more Cre+ cells in the CeM than CeL. The inferior performance of the Con/Foff virus may have resulted from incomplete coverage of Flp expression in CeL neurons. Cre-positive CeL cells that do not co-express Flp would express the Con/Foff reporter and thereby lower the CeM:CeL ratio.

The CeL is largely composed of three cell populations: CeL^PKCδ^ neurons that inhibit food or water consumption (this study and ^6,7,55–57^), and Sst+ neurons that can be further subdivided into CeL^Sst^ and CeL^Nts/Tac2^ cells ^7, 13, 21^. Both Sst+ populations in CeL heavily overlap with Htr2a-Cre expressing cells ^13^. Here, by targeting Sst-Cre- or Htr2a-Cre-positive cells in the CeL, we obtained no evidence that they could promote water or food consumption. This was a surprising finding in light of earlier reports showing that Sst+, Nts+, Pnoc+, and Htr2a+ neurons promoted water, palatable fluid, and/or palatable food uptake ^5, 7, 22, 23^. However, these reports did not distinguish between CeL and CeM subpopulations, raising the possibility that it was indeed the CeM subpopulations that drove the behavior. The study by Kim et al., (2017) ^7^, attempted to target the anatomical subregions with stereotaxic viral injections and reported that both CeL and CeM subpopulations of Sst+ and Nts/Tac2+ neurons promoted water uptake. This result is contrary to ours, and may in part be due to the different techniques employed. Further experiments would be necessary to solve this discrepancy.

Contrary to the CeL, the Sst+ and Htr2a-Cre+ subpopulations in the CeM are largely separate populations ^13^. Here, we show that CeM^Sst^ neurons promote water, but not solid food uptake, whereas CeM^Htr2a^ neurons promote water and food uptake. These findings are consistent with and extend earlier observations: Nts+ neurons (a subpopulation of Sst neurons) promote ethanol and palatable fluid, but not solid food, consumption ^23^; photoactivation of Tac2+ or CRH+ cells (subpopulations of Sst neurons) has no positive effect on feeding ^6^; Pnoc+ neurons (heavily overlapping with Htr2a-Cre-expressing neurons) promote palatable food consumption ^22^.

The optogenetic experiments are supported by *in vivo* calcium imaging data which indicate that the majority of CeM^Htr2a^ and CeM^Sst^ neurons increased their activities during consumption of water and food rewards. CEBRA analysis revealed that their activity dynamics contributed to and decoded specific behavioral features. Depending on the type of rewards, CeM^Htr2a^ and CeM^Sst^ neurons were recruited into different ensembles whose activities correlated positively or negatively with reward consumption. The highest positive correlation of CeM^Htr2a^ neurons was with feeding (59%), consistent with their observed function in food consumption. When switching between rewards of different physical attributes (e.g. solid food and liquid Fresubin), CeM^Htr2a^ neurons more often displayed a stable correlation, while CeM^Sst^ neurons more often changed their activities. These results are consistent with previously suggested models ^11, 12^ in which CeA^Sst^ neurons participate in discriminating between stimuli that differ in their sensory/physical properties, such as taste and texture. In contrast to CeM^Sst^ neurons, CeM^Htr2a^ neurons may encode a broader range of information, such as the innately affective properties of the stimulus and the animal’s internal hunger state. CeM^Htr2a^ cells are activated by fasting and the hunger hormone ghrelin ^13^ and here, we found that a higher percentage of CeM^Htr2a^ cells were positively correlated with feeding when the animals were hungry compared to when they were fed and consuming Fresubin. Hence, CeM^Htr2a^ neuron activity is less correlated with the physical attributes of the stimulus, and more with its palatability and rewarding properties.

If the activities of CeM^Sst^ neurons increase during water licking and feeding bouts, why is ectopic activation of these neurons sufficient to promote drinking, but not food consumption? Previously, it was shown that the activities of Sst+ neurons (in the entire CeA) arose later than the animal’s licking responses following water delivery, suggesting that the activities of these neurons did not promote licking ^12^. Instead, Sst+ neurons may drive water consumption by conveying stimulus-specific signals to downstream reward centers. Solid food consumption involves additional aspects such as handling and biting the food, and these aspects may only be driven by CeM^Htr2a^, but not CeM^Sst^ neurons. Further work will be necessary to test this hypothesis.

At the population level, Sst+ CeA neurons were previously shown to encode a range of appetitive and aversive stimuli. Many individual Sst+ neurons displayed high selectivity to only one class of stimuli (‘specializers’), some responded to multiple stimuli, sometimes of opposite valence (‘generalizers’) ^12^. Similar heterogeneity may be present in other CeA populations, such as Nts+ neurons which promote the uptake of palatable fluids, but not solid food ^23^. Here, we also recorded the activities of individual CeM neurons when animals experienced a switch between an appetitive and an aversive liquid. For CeM^Htr2a^ neurons, the fraction of specializers was significantly larger than the fraction of generalizers when switching from saccharin to quinine solution. Interestingly, for the same switch in stimuli, the pattern was opposite for CeM^Sst^ cells, with generalizers outnumbering specializers. We conclude that the CeM^Htr2a^ population contains more cells that innately specialize in encoding valence-specific stimuli than the CeM^Sst^ population. In other words, for CeM^Htr2a^ neurons that may integrate physical and rewarding properties of the stimulus with the animal’s hunger state, the switch from saccharin to quinine represents an important valence switch that causes a change in activity for most neurons. For CeM^Sst^ cells that mainly respond to stimuli that differ in their sensory/physical properties, a switch between a sweet and a bitter liquid represents a minor difference in physical attributes that did not cause a change in activity for most neurons.

Previously, the activities of Sst+ and Htr2a-Cre-expressing neurons (of the entire CeA) were shown to be intrinsically rewarding in RTPP assays, and intrinsically reinforcing in intracranial self-stimulation and reward learning assays ^5, 7, 42^. Here, we show that both CeM subtypes drive RTPP and conditioned flavor preference. It is likely that this activity contributes to the promotion of water or food uptake. The CeL subtypes of Sst+ and Htr2a-Cre-expressing neurons also displayed modest reinforcing activities in the conditioned flavor preference assay, but not in RTPP. These observations are in line with the notion that Sst+ neurons are required for learning, that Sst+ neurons projecting to substantia nigra (SN) dopaminergic neurons participate specifically in reward learning ^12^, and that both CeL and CeM subtypes form projections to the SN (this study).

The appetitive inputs that lead to activation of CeM subtypes are not well understood. Brain regions controlling water intake include the anterior cingulate cortex and their projections to the amygdala complex ^58^ and the peri-locus coeruleus ^59^. Food-related inputs may come from insular cortex ^20, 60^, PBN ^45, 49^, the arcuate nucleus ^5, 61^ and the parasubthalamic nucleus ^5, 62^, from fasting, and the hunger hormone ghrelin^13^. Here, we have shown that both CeM subtypes are under inhibitory control of CeA^PKCδ^ neurons residing in the CeL. When animals reach satiety during consumption of water or food, the behaviour is terminated by activation of CGRP+ PBN neurons ^45, 48^ that project to the CeA and activate PKCδ neurons. Conversely, when animals are thirsty or hungry, CeA^PKCδ^ neurons may become inhibited, either through the local CeL inhibitory network, or by long-range inhibitory inputs to CeA^PKCδ^ neurons, and this will disinhibit CeM^Htr2a^ and CeM^Sst^ neurons favoring the expression of appetitive behavior.

Major long-range projections of CeM subtypes to other brain regions involved in rewarding and consummatory behaviors include the BNST, IPAC, LH, SN, and PBN ^5, 20, 23, 41–43^. The PBN is an interesting output region because it receives projections from CeM^Htr2a^ neurons driving food consumption, from CeM^Sst^ neurons driving water consumption, and from CeM^Dlk1^ neurons suppressing food intake during nausea ^63^. It is likely that CeM^Htr2a^→PBN and CeM^Sst^→PBN projectors target different microcircuits in the PBN, since their optogenetic activation elicited different behaviors. Anorexigenic CeM^Dlk1^→PBN projectors may interfere with the appetitive CeM→PBN projectors, for example, through a presynaptic inhibition mechanism. Future work is needed to work out the information flow from the CeM to the PBN.

In conclusion, the present manuscript provides a detailed functional characterization of molecularly- and anatomically-defined appetitive CeA subpopulations. Our findings indicate that neurons driving food or water consumption are confined to the CeM and that separate CeM subpopulations regulate water only (CeM^Sst^), versus water or food consumption (CeM^Htr2a^). The response properties of these CeM neurons show interesting differences regarding reward value and physical attributes of the stimuli. In the future, further characterization of these circuits, including modulation by neuropeptides and comparative evolutionary studies in healthy and diseased subjects may provide insights into the etiology of eating and drinking disorders and may help to develop therapeutic strategies to combat these common problems.

## ACKNOWLEDGEMENTS

We thank Soo Jin Min-Weissenhorn and the Transgenic Service for help with generating transgenic mouse lines, Jens F. Rehfeld, University of Copenhagen, for providing the Wfs1 antibody, and Minh Chau Mai and Emily Russell for help with management of the animal colonies. This study was supported by the Max-Planck Society and the European Research Council under the European Union’s Horizon 2020 research and innovation programme (No. 885192).

## AUTHOR CONTRIBUTIONS

FF and RK conceptualized the study and designed experiments. FF performed most of the experiments. SC performed the CEBRA analysis. CP performed and analyzed electrophysiology experiments. LG assisted with histology, immunohistochemistry, microscopy, and image processing. PLAM generated the Wfs1-FlpoER transgene and participated in the design of the PKCδ-Flpo transgene and in establishing both transgenic mouse lines. CR and KD provided intersectional viruses. FF and RK wrote the manuscript with input from all authors. RK supervised and provided funding.

## Methods

### Animals

Experiments were always performed using adult mice (> 12 weeks). The Htr2a-Cre BAC transgenic line (stock Tg(Htr2a-Cre)KM208Gsat/Mmucd) was imported from the Mutant Mouse Regional Resource Center 482 (https://www.mmrrc.org/). SOM-IRES-cre (SSTtm2.1(cre)Zjh/J) mice were acquired from the Jackson Laboratory. Ai9lsl−TdTomato (B6.Cg-Gt(ROSA) 26Sortm9(CAG-tdTomato)Hze/J)^64^, FPDI (B6;129S6Gt(ROSA)26Sortm9 (CAG-mCherry, -CHRM4*)Dym)^34^ mouse lines were as described previously. All mice were backcrossed into a C57BL/6NRj background (Janvier Labs - http://www.janvier-labs.com). Both male and female mice were used and all the experiments were performed following regulations from the government of Upper Bavaria.

### Generation of Wfs1-FlpoER transgenic mice

For the generation of Wfs1-FlpoER mice, a BAC homologous recombination method was used. The BAC clone RP23-405O19 (CHORI) was targeted with a FLPoERT2 expression cassette^65^ and a WPRE sequence followed by the bovine growth hormone polyadenylation signal (pA) and a kanamycin resistance cassette. The BAC homologous recombination cassette was assembled in the pcDNA3.1 (+) vector (Invitrogen) as follows: Homology arms A and B were designed to flank mouse Wfs1 exon 2. A construct containing the homology arm A and the NLSFlpoERT2WPREpA sequences was synthetized by Eurofins and cloned into the pUC57 vector (GenScript). Homology arm B was synthetized by PCR using the following primer sets: 5’-CGATATCAACTCAGGCACC -3’ (forward primer for arm B), 5’-AATCTCGAGCAGGGACACTG -3’ (reverse primer for arm B). First, pCDNA3.1 (+) was digested with NheI/AflII to insert the homologous arm A – NLSFlpoERT2WPREpA fragment into this site. Second, the kanamycin resistance cassette (Gene Bridges GmbH) was cloned into the AflII/EcoRV site before the arm B. Then, the homologous arm B was inserted into the EcoRV/XhoI site. This targeting vector was digested with NheI/XhoI followed by purification of the insert on a 0.7% agarose gel. Homologous recombinant BACs were obtained using established methods^66, 67^, screened by PCR and verified by Southern blotting. Modified BAC DNA was prepared using the large construct DNA purification kit - NucleoBond^®^ Xtra BAC (Macherey-Nagel), linearized with PmeI and purified through a Sepharose™ separation column^68^. BAC DNA (2.2 ng/μl) was injected into pronuclei of fertilized oocytes of C57BL/6 mice. BAC transgenic mice were identified by PCR using the primers 5’ GCTCTATTCAGGACATTTTCACATCTCTAC 3’ and 5’ CCTCTCGAATCTCTCCACGAAC 3’

### Generation of PKCδ-Flpo transgenic mice

A recombineering protocol was used to insert a NLSFLPo-pA expression cassette^69^ into the ATG start site of the PKCdelta locus on the BAC clone RP23-283B12 (CHORI). The wild-type loxP site present in the RP23 BAC backbone was replaced by a piggyBAC-ampR cassette. Plasmid construction and BAC modification was done by Gene Bridges GmbH. BAC DNA for pronuclear injection was prepared using the large construct DNA purification kit - NucleoBond^®^ Xtra BAC (Macherey-Nagel), linearized with PI-SceI and purified over a home-made Sepharose CL4B column (Johansson et al., 2010). Fractions were analyzed by pulsed-field gel electrophoresis to identify the sample with the highest concentration of linearized BAC DNA and lowest concentration of vector DNA. Linearized BAC DNA was injected at a concentration of 3.65 ng/μl into pronuclei of fertilized oocytes of C57BL/6 mice. BAC transgenic mice were identified by PCR using the primers 5’ AAACTGCATCACCTTCTCACATCTCC 3’ and 5’ CTCTCGAATCTCTCCACGAACTGC 3’.

### Viral constructs

The following AAV viruses were purchased from the University of North Carolina Vector Core (https://www.med.unc.edu/genetherapy/vectorcore: AAV5-Ef1a-DIO-eNpHR3.0-mCherry, AAV5-Ef1a-DIO-mCherry, AAV5-Ef1a-DIO-hChR2(H134R)-EYFP-WPRE-pA, AAV5-Ef1a-DIO-EYFP-WPRE-pA, AAV5-hSyn-Con/Fon-hChR2(H134R)-EYFP-WPRE, AAV5-hSyn-Con/Fon-EYFP-WPRE, AAV5-hSyn-Con/Foff-hChR2(H134R)-EYFP-WPRE, AAV5-hSyn-Con/Foff-EYFP-WPRE, AAV5-EF1a-fDIO-hChR2(H134R)-EYFP-WPRE.

AAV8-nEF-Con/Foff iC++-EYFP, AAV8-nEF-Con/Foff–EYFP and AAV8-EF1a-Con/Foff–GCaMP6m viruses were generated as described ^31^.

### Viral injections

Mice were anaesthetized using isoflurane (Cp-pharma) and placed on a heating pad on a stereotaxic frame (Model 1900 – Kopf Instruments). Carprofen (Rimadyl – Zoetis) (5 mg/kg body weight) was given via subcutaneous injection. Mice were bilaterally (or unilaterally for calcium imaging experiments) injected using glass pipettes (#708707, BLAUBRAND intraMARK) with 0.3 µl of virus in the CeA by using the following coordinates calculated with respect to bregma: for the CeA and CeL: −1.20 mm anteroposterior, ± 2.87 mm lateral, −4.65 to -4.72 mm ventral; for the CeM −1.155 mm anteroposterior, ± 2.87 mm lateral, −4.65 to -4.72 mm ventral. The following stereotaxic coordinates were used for the PBN: -5.2 mm anteroposterior, ±1.4 mm lateral, -3.85 mm ventral. Virus was allowed to be expressed for a minimum duration of 3 weeks before histology or behavioral paradigms. For animals not undergoing implant surgery, the incision was sutured.

### Optic fiber implants

Mice used in optogenetic experiments were implanted with optic fibers (200-µm core, 0.22 NA, 1.25-mm ferrule - Thorlabs) above the CeA (−4.35 mm ventral from bregma) or the PBN (-3.6 mm ventral from bregma) immediately after viral injection. The skull was first protected with a layer of histo glue (Histoacryl, Braun), the fibers were then fixed to the skull using UV light-curable glue (Loctite AA3491 - Henkel), and the exposed skull was covered with dental acrylic (Paladur - Heraeus).

### GRIN lens implantation and baseplate fixation

Three weeks after GCaMP6s viral injection in the CeA, mice were implanted with a gradient index (GRIN) lens. At the same coordinates of the injection, a small craniotomy was made and a 20G needle was slowly lowered into the brain to clear the path for the lens to a depth of -4.5 mm from bregma. After retraction of the needle, a GRIN lens (ProView lens; diameter, 0.5 mm; length, ∼8.4 mm, Inscopix) was slowly implanted above the CeA and then fixed to the skull using UV light-curable glue (Loctite AA3491 - Henkel). The skull was first protected with histo glue (Histoacryl, Braun), and the implant fixed with dental acrylic (Paladur - Heraeus). 4 to 8 weeks after GRIN lens implantation, mice were “baseplated” under anesthesia. Briefly, in the stereotaxic setup, a baseplate (BPL-2; Inscopix) was positioned above the GRIN lens, adjusting the distance and the focal plane until the neurons were visible. The baseplate was fixed using C&B Metabond (Parkell). A baseplate cap (BCP-2, Inscopix) was left in place to protect the lens.

### Tamoxifen

Tamoxifen was prepared by dissolving 20mg tamoxifen in 100% ethanol to obtain a final concentration of 40 mg/ml. The solution was then diluted 1:1 with Kolliphor ® EL (Sigma). By heating and stirring the mixture, the ethanol evaporates. After that, 100 µl aliquots were frozen until use. Before use, the stock solution was diluted with PBS to the desired concentration and the pH value of 7.4 was confirmed. Mice were injected with approximately 200 μl of tamoxifen solution (200 mg per kilogram of body weight) for three days every other day. The mice that underwent brain viral injection received the first injection immediately after the surgery.

### Behavioral assays

Mice were bilaterally tethered to optic fiber patch cables (Doric Lenses or Thorlabs) via a mating sleeve (Thorlabs). The patch cables were connected via a rotary joint (Doric Lenses) to a 473nm or 561nm (CNI lasers) laser. Photoactivation and photoinhibition experiments were conducted with 10-15mW 10ms, 473nm light pulses at 20 Hz, using a pulser (Prizmatix) controlled by the Ethovision software XT 14 (Noldus), or 561nm 15mW constant light.

Drinking behavior: For water deprivation experiments, mice were water deprived and the following day trained to drink in a 32 cm x 35 cm plastic arena for 30 minutes. This training was mainly done to habituate the mice to the new setup and train them to drink from specific pipettes. For photoactivation experiments, mice were then tested the following two days in the setup with access to two pipettes of water with a sequence of 10 min laser OFF/10 min laser ON (10 min laser ON/10 min laser OFF for PKCδ mice), or simply for 30 min laser ON. The two experiments were performed in a randomized order. For experiments in normal conditions, the same protocols were used, but the mice had *ad libitum* water in their cage. For photoinhibition experiments, water-deprived mice had access to water for 30 min while photoinhibited the day after the training. The amount of water was manually measured.

Feeding behavior: The experiments were conducted in a 32 cm x 35 cm plastic arena containing two plastic cups in opposite corners, one with dustless precision pre-weighed food pellets (20 mg each) (Bio-Serv-F0071) and one empty. Experiments were conducted over 40 min sessions and the remaining food was weighed. The day before the experiment, mice were habituated to the new food by placing some of these pellets into the cage together with the normal food.

Palatable reward consumption: Mice were food deprived for 16 hours and allowed to consume a palatable reward solution (Fresubin, 2kcal/ml) for 30-45 min. The mice went back to *ad libitum* food and the following day were tested for 30 min for Fresubin consumption while photoinhibited.

Real-time place preference: ChR2-expressing mice and corresponding controls were allowed to freely navigate in a custom-made plexiglas two-chambered arena (50x25x25cm) for 20 min. ChR2-expressing mice and controls received 20Hz 473nm photostimulation in one compartment. The experiment was repeated for two consecutive days alternating the photostimulated and neutral chambers.

Conditioned flavor preference: Mice deprived of water deprived mice for two consecutive days were given the choice between two nonnutritive flavored liquids (0.3 % grape or cherry and 0.15% saccharin) for 30 min. The preferred taste was defined as the taste from the two that the mice drank more of. Conditioning was conducted over four consecutive days with two sessions per day. The least preferred taste was paired with optogenetic activation. In conditioning session one, the least preferred taste was paired with light for 15 minutes. In conditioning session two, the mice were presented with the more preferred taste in the absence of photostimulation. The order of the sessions was inverted each day, occurring 4-6 hours apart. Conditioned flavor preference was tested the day after the final conditioning session, when the mice were presented with both tastes.

Open field: Mice were allowed to explore a custom-built plexiglas arena (40 cm×40 cm×25 cm) for 10 min. During the whole experiment, mice received photoactivation or photoinhibition.

### *In vivo* calcium imaging of freely moving mice

All *in vivo* imaging experiments were conducted on freely moving mice. GCaMP6m fluorescence signals were acquired using a miniature integrated fluorescence microscope system (nVoke – Inscopix) secured in the baseplate holder before each imaging session. Mice were habituated to the miniscope procedure for 3 days before behavioral experiments for 30 min per day. Settings were kept constant within subjects and across imaging sessions. Image acquisition and behavior were synchronized using the data acquisition box of the nVoke Imaging System (Inscopix) triggered by the Ethovision XT 14 software (Noldus) through a TTL box (Noldus) connected to the USB-IO box from the Ethovision system (Noldus). For drinking experiments, mice were water deprived and trained the next day to drink in a 32 cm x 35 cm plastic arena for 30 minutes. Calcium activity was recorded the following day during a 10-minute period of habituation without water and a subsequent 10-minute period of exposure to water. For feeding experiments, mice were food restricted and trained the following day to consume dustless precision pre-weighed food pellets (20 mg each) (Bio-Serv-F0071) in a 32 cm x 35 cm plastic arena containing two plastic cups in opposite corners, one of which empty. The next day, after 10 min of recording without food, animals were exposed to food for an additional 10 minutes and their calcium activity was recorded. The Fresubin experiment was conducted as previously described in this manuscript. On the test day, the calcium activity of fed mice was recorded during a 10-minute period of habituation followed by a 10-minute period after the introduction of Fresubin. To study water/saccharin/quinine behavior, mice were water deprived and habituated the following day to water administration in the experimental setup. On the next day, the experiment was carried out in three consecutive 10-minute sessions. During the first, second and third session, water, saccharin (0.15%) and quinine (10mM) were respectively administered at minutes 1 and 6. For imaging data processing and analysis we used IDPS (Inscopix data 721 processing software) version 1.8.0.

### CEBRA analysis

To understand if the recorded behaviors contributed to changes in neural activity, we applied the CEBRA pipeline for feature selection. If certain behaviors account significantly for neural activity, we would have expected to see clear structures in the CEBRA embeddings compared to shuffled neural data. In brief, we chose the CEBRA Hybrid model (self-supervised) with parameters used in this study as below:

Hybrid model, model_architecture= ‘offset5-model-mse’, conditional= “time-delta”, hybrid = True, distance= ‘euclidean’, batch_size= 600-1200, learning_rate=1e-4. Iteration = 8000.

To examine the association between CEBRA embeddings and behaviors we applied the Random Forest (RF) classifier for behavioral decoding. In brief, we fitted behavioral labels with our CEBRA embeddings, we used 70% of the data as training data and 30% of the data as test data (with data splitting in random orders). In addition, we permuted the data 1000 times randomly and tested it with the trained RF classifier. We inspected the model by checking the decoding accuracy and the out of bag (OOB) error. To prevent overfitting of the model, we decoded behavioral labels first within the same animal (Fig.5 f,g,i,j ; Extended Data Fig.5 g,h; Extended Data Fig.6 d,e), then we decoded the behavioral labels across animals (Fig.5 f,g ; Extended Data Fig.5 g).

### Brain-Slice preparation and electrophysiological recordings

Mice were anesthetized using isoflurane and subsequently sacrificed by decapitation. The brains were then placed in a cutting solution composed of 95% oxygen and 5% carbon dioxide, with a mixture of 30 mM NaCl, 4.5 mM KCl, 1 mM MgCl_2_, 26 mM NaHCO_3_, 1.2 mM NaH_2_PO_4_, 10 mM glucose, and 194 mM sucrose. The brains were sliced to a thickness of 280 μm using a Leica VT1000S vibratome (Germany) and transferred to an artificial cerebrospinal fluid (aCSF) solution containing 124 mM NaCl, 4.5 mM KCl, 1 mM MgCl_2_, 26 mM NaHCO_3_, 1.2 mM NaH_2_PO_4_, 10 mM glucose, and 2 mM CaCl_2_ (310-320 mOsm), saturated with 95% O_2_/5% CO_2_ at 32°C for 1 hour before being brought to room temperature. The brain slices were then placed in a recording chamber that was continuously perfused with aCSF solution, also saturated with 95% O_2_/5% CO_2_, at a temperature of 30-32°C.

Whole-cell patch-clamp recordings were carried out as previously described^20^. Patch pipettes were fabricated from filament-containing borosilicate micropipettes (World Precision Instruments) using a P-1000 micropipette puller (Sutter Instruments, Novato, CA) to achieve a resistance of 6-8 MΩ. The intracellular solution consisted of 130 mM potassium gluconate, 10 mM KCl, 2 mM MgCl_2_, 10 mM HEPES, 2 mM Na-ATP, 0.2 mM Na_2_GTP at a pH of 7.35, and an osmotic pressure of 290 mOsm. The brain slices were visualized using a fluorescence microscope equipped with IR-DIC optics (Olympus BX51). The holding potential for excitatory postsynaptic currents (EPSC) was set at -70 mV and 0 mV for inhibitory postsynaptic currents (IPSC). Data was acquired using a MultiClamp 700B amplifier, Digidata 1550 digitizer (Molecular Devices), and analyzed using Clampex 10.3 software (Molecular Devices, Sunnyvale, CA). The data was sampled at 10 kHz and filtered at 2 kHz, and further analyzed using Clampfit (Molecular Devices).

For optogenetic studies, neurons were stimulated through the use of a multi-LED array system (CoolLED) connected to an Olympus BX51 microscope.

### Histology

Mice were anesthetized with ketamine/xylazine solution (Medistar and Serumwerk) (100 mg/kg and 16 mg/kg, respectively) and transcardially perfused with phosphate-buffered saline (PBS), followed by 4% paraformaldehyde (PFA) (1004005, Merck) (w/v) in PBS. Extracted brains were post-fixed overnight at 4°C in 4% PFA (w/v) in PBS, embedded in 6% agarose and sliced using a Vibratome (VT1000S - Leica) into 50-μm free-floating coronal sections.

### Immunohistochemistry

Brain sections were blocked at room temperature for two hours in 5% donkey serum (Biozol JIM-017-000-121) diluted in 1X PBS 0.5% TritonX-100 and then incubated with primary antibody at 4°C overnight in the same solution. Primary antibodies: goat anti-mCherry (1:1000) (Origene AB0040-200), chicken anti-GFP (1:500) (Aves, GFP-1020), rabbit anti-Wfs1 (1:200) (made by the lab of Prof. Dr. Jens F. Rehfeld, University of Copenhagen, Denmark), mouse anti-PKCδ (1:100; 610398, BD Biosciences). After incubation with primary antibody, the sections were washed in 1X PBS (3x 10 minutes) and incubated in secondary antibody in 1X PBS 0.5% TritonX-100 at 4°C overnight. Secondary antibodies: Alexa Fluor donkey anti-rabbit/goat/chicken 488/Cy3/647, (1:300) (Jackson). After 3x10 min washes in 1X PBS, sections were incubated in DAPI and coverslipped (Dako).

### Microscopy and image processing

A Leica SP8 confocal microscope equipped with a 20×/0.75 IMM or 10x/0.30 FLUOTAR objective (Leica) was used to acquire Fluorescence z-stack images. Full views of brain slices were acquired using the tile scan and automated mosaic merge functions of Leica LAS X software. Images were minimally processed with ImageJ software (NIH) to adjust for brightness and contrast for optimal representation of the data. For all quantifications of brain sections, ImageJ was used to manually count the cells.

### Statistical analysis

No statistical methods were used to pre-determine sample sizes. The numbers of samples in each group were based on those in previously published studies. Statistical analyses were performed with Prism 9 (GraphPad) and all statistics are indicated in the figure legends. T-tests or two-way ANOVA with Bonferroni post-hoc tests were used for individual comparisons of normally distributed data. Normality was assessed using Shapiro-Wilk test. When normality was not assumed, Mann-Whitney U test and Wilcoxon signed-rank test were performed for individual comparisons. After the conclusion of experiments, virus-expression and implants placement were verified. Mice with very low or null virus expression were excluded from analysis.

## REFERENCES

1. Balleine, B.W. & Killcross, S. Parallel incentive processing: an integrated view of amygdala function. Trends Neurosci 29, 272–279 (2006).

2. Janak, P.H. & Tye, K.M. From circuits to behaviour in the amygdala. Nature 517, 284–292 (2015).

3. Davis, M. & Whalen, P.J. The amygdala: vigilance and emotion. Mol Psychiatry 6, 13–34 (2001).

4. Everitt, B.J., Cardinal, R.N., Parkinson, J.A. & Robbins, T.W. Appetitive behavior: impact of amygdala-dependent mechanisms of emotional learning. Ann N Y Acad Sci 985, 233–250 (2003).

5. Douglass, A.M., et al. Central amygdala circuits modulate food consumption through a positive-valence mechanism. Nat Neurosci 20, 1384–1394 (2017).

6. Cai, H., Haubensak, W., Anthony, T.E. & Anderson, D.J. Central amygdala PKC-δ(+) neurons mediate the influence of multiple anorexigenic signals. Nat Neurosci 17, 1240–1248 (2014).

7. Kim, J., Zhang, X., Muralidhar, S., LeBlanc, S.A. & Tonegawa, S. Basolateral to Central Amygdala Neural Circuits for Appetitive Behaviors. Neuron 93, 1464–1479.e1465 (2017).

8. Morales, I. & Berridge, K.C. ’Liking’ and ’wanting’ in eating and food reward: Brain mechanisms and clinical implications. Physiol Behav 227, 113152 (2020).

9. Chen, G., et al. Distinct reward processing by subregions of the nucleus accumbens. Cell Rep 42, 112069 (2023).

10. Kargl, D., et al. The amygdala instructs insular feedback for affective learning. Elife 9 (2020).

11. Ponserre, M., Fermani, F., Gaitanos, L. & Klein, R. Encoding of Environmental Cues in Central Amygdala Neurons during Foraging. The Journal of Neuroscience 42, 3783–3796 (2022).

12. Yang, T., et al. Plastic and stimulus-specific coding of salient events in the central amygdala. Nature 616, 510–519 (2023).

13. Peters, C., et al. Transcriptomics reveals amygdala neuron regulation by fasting and ghrelin thereby promoting feeding. Sci Adv 9, eadf6521 (2023).

14. Livneh, Y., et al. Homeostatic circuits selectively gate food cue responses in insular cortex. Nature 546, 611–616 (2017).

15. Fadok, J.P., Markovic, M., Tovote, P. & Lüthi, A. New perspectives on central amygdala function. Curr Opin Neurobiol 49, 141–147 (2018).

16. Han, W., et al. Integrated Control of Predatory Hunting by the Central Nucleus of the Amygdala. Cell 168, 311–324.e318 (2017).

17. Mike, J.F.R., Shelley, M.W. & Kent, C.B. Optogenetic Excitation of Central Amygdala Amplifies and Narrows Incentive Motivation to Pursue One Reward Above Another. The Journal of Neuroscience 34, 16567 (2014).

18. Sevil, D., Daniela, P. & Denis, P. Central Amygdala Activity during Fear Conditioning. The Journal of Neuroscience 31, 289 (2011).

19. LeDoux, J.E., Iwata, J., Cicchetti, P. & Reis, D.J. Different projections of the central amygdaloid nucleus mediate autonomic and behavioral correlates of conditioned fear. J Neurosci 8, 2517–2529 (1988).

20. Ponserre, M., Peters, C., Fermani, F., Conzelmann, K.K. & Klein, R. The Insula Cortex Contacts Distinct Output Streams of the Central Amygdala. J Neurosci 40, 8870–8882 (2020).

21. Wang, Y., et al. Multimodal mapping of cell types and projections in the central nucleus of the amygdala. Elife 12 (2023).

22. Hardaway, J.A., et al. Central Amygdala Prepronociceptin-Expressing Neurons Mediate Palatable Food Consumption and Reward. Neuron 102, 1037–1052.e1037 (2019).

23. Torruella-Suárez, M.L., et al. Manipulations of Central Amygdala Neurotensin Neurons Alter the Consumption of Ethanol and Sweet Fluids in Mice. J Neurosci 40, 632–647 (2020).

24. McCullough, K.M., Daskalakis, N.P., Gafford, G., Morrison, F.G. & Ressler, K.J. Cell-type-specific interrogation of CeA Drd2 neurons to identify targets for pharmacological modulation of fear extinction. Transl Psychiatry 8, 164 (2018).

25. Dilly, G.A., Kittleman, C.W., Kerr, T.M., Messing, R.O. & Mayfield, R.D. Cell-type specific changes in PKC-delta neurons of the central amygdala during alcohol withdrawal. Translational Psychiatry 12, 289 (2022).

26. Yu, K., Garcia da Silva, P., Albeanu, D.F. & Li, B. Central Amygdala Somatostatin Neurons Gate Passive and Active Defensive Behaviors. J Neurosci 36, 6488–6496 (2016).

27. Wilson, T.D., et al. Dual and Opposing Functions of the Central Amygdala in the Modulation of Pain. Cell Rep 29, 332–346.e335 (2019).

28. Fadok, J.P., et al. A competitive inhibitory circuit for selection of active and passive fear responses. Nature 542, 96–100 (2017).

29. Madisen, L., et al. Transgenic mice for intersectional targeting of neural sensors and effectors with high specificity and performance. Neuron 85, 942–958 (2015).

30. Fenno, L.E., et al. Targeting cells with single vectors using multiple-feature Boolean logic. Nat Methods 11, 763–772 (2014).

31. Fenno, L.E., et al. Comprehensive Dual-and Triple-Feature Intersectional Single-Vector Delivery of Diverse Functional Payloads to Cells of Behaving Mammals. Neuron 107, 836–853.e811 (2020).

32. Becker, J.A., et al. Transcriptome analysis identifies genes with enriched expression in the mouse central extended amygdala. Neuroscience 156, 950–965 (2008).

33. Luuk, H., et al. Distribution of Wfs1 protein in the central nervous system of the mouse and its relation to clinical symptoms of the Wolfram syndrome. J Comp Neurol 509, 642–660 (2008).

34. Ray, R.S., et al. Impaired respiratory and body temperature control upon acute serotonergic neuron inhibition. Science 333, 637–642 (2011).

35. Berndt, A., et al. Structural foundations of optogenetics: Determinants of channelrhodopsin ion selectivity. Proc Natl Acad Sci U S A 113, 822–829 (2016).

36. Peters, C., et al. Transcriptomics reveals amygdala neuron regulation by fasting and ghrelin thereby promoting feeding. bioRxiv, 2022.2010.2021.513224 (2022).

37. Schneider, S., Lee, J.H. & Mathis, M.W. Learnable latent embeddings for joint behavioural and neural analysis. Nature 617, 360–368 (2023).

38. Ch’ng, S., Fu, J., Brown, R.M., McDougall, S.J. & Lawrence, A.J. The intersection of stress and reward: BNST modulation of aversive and appetitive states. Progress in Neuro-Psychopharmacology and Biological Psychiatry 87, 108–125 (2018).

39. Jennings, J.H., Rizzi, G., Stamatakis, A.M., Ung, R.L. & Stuber, G.D. The inhibitory circuit architecture of the lateral hypothalamus orchestrates feeding. Science 341, 1517–1521 (2013).

40. Furlan, A., et al. Neurotensin neurons in the extended amygdala control dietary choice and energy homeostasis. Nat Neurosci 25, 1470–1480 (2022).

41. Lee, H.J., et al. Role of amygdalo-nigral circuitry in conditioning of a visual stimulus paired with food. J Neurosci 25, 3881–3888 (2005).

42. Steinberg, E.E., et al. Amygdala-Midbrain Connections Modulate Appetitive and Aversive Learning. Neuron 106, 1026–1043.e1029 (2020).

43. Ryan, P.J., Ross, S.I., Campos, C.A., Derkach, V.A. & Palmiter, R.D. Oxytocin-receptor-expressing neurons in the parabrachial nucleus regulate fluid intake. Nat Neurosci 20, 1722–1733 (2017).

44. Bloodgood, D.W., et al. Kappa opioid receptor and dynorphin signaling in the central amygdala regulates alcohol intake. Mol Psychiatry 26, 2187–2199 (2021).

45. Campos, C.A., Bowen, A.J., Schwartz, M.W. & Palmiter, R.D. Parabrachial CGRP Neurons Control Meal Termination. Cell Metab 23, 811–820 (2016).

46. Han, S., Soleiman, M.T., Soden, M.E., Zweifel, L.S. & Palmiter, R.D. Elucidating an Affective Pain Circuit that Creates a Threat Memory. Cell 162, 363–374 (2015).

47. Palmiter, R.D. The Parabrachial Nucleus: CGRP Neurons Function as a General Alarm. Trends Neurosci 41, 280–293 (2018).

48. Campos, C.A., Bowen, A.J., Roman, C.W. & Palmiter, R.D. Encoding of danger by parabrachial CGRP neurons. Nature 555, 617–622 (2018).

49. Carter, M.E., Soden, M.E., Zweifel, L.S. & Palmiter, R.D. Genetic identification of a neural circuit that suppresses appetite. Nature 503, 111–114 (2013).

50. Pauli, J.L., et al. Molecular and anatomical characterization of parabrachial neurons and their axonal projections. eLife 11, e81868 (2022).

51. Michael, C.C., et al. Parabrachial Complex: A Hub for Pain and Aversion. The Journal of Neuroscience 39, 8225 (2019).

52. Biglari, N., et al. Functionally distinct POMC-expressing neuron subpopulations in hypothalamus revealed by intersectional targeting. Nat Neurosci 24, 913–929 (2021).

53. Poulin, J.F., et al. Mapping projections of molecularly defined dopamine neuron subtypes using intersectional genetic approaches. Nat Neurosci 21, 1260–1271 (2018).

54. Britz, O., et al. A genetically defined asymmetry underlies the inhibitory control of flexor– extensor locomotor movements. eLife 4, e04718 (2015).

55. Sanchez, M.R., et al. Dissecting a disynaptic central amygdala-parasubthalamic nucleus neural circuit that mediates cholecystokinin-induced eating suppression. Molecular Metabolism 58, 101443 (2022).

56. Schnapp, W.I., et al. Development of activity-based anorexia requires PKC-δ neurons in two central extended amygdala nuclei. Cell Rep 43, 113933 (2024).

57. Park-York, M., Boghossian, S., Oh, H. & York, D.A. PKCθ expression in the amygdala regulates insulin signaling, food intake and body weight. Obesity 21, 755–764 (2013).

58. Zhao, Z., et al. A Novel Cortical Mechanism for Top-Down Control of Water Intake. Curr Biol 30, 4789–4798.e4784 (2020).

59. Gong, R., Xu, S., Hermundstad, A., Yu, Y. & Sternson, S.M. Hindbrain Double-Negative Feedback Mediates Palatability-Guided Food and Water Consumption. Cell 182, 1589–1605.e1522 (2020).

60. Zhang-Molina, C., Schmit, M.B. & Cai, H. Neural Circuit Mechanism Underlying the Feeding Controlled by Insula-Central Amygdala Pathway. iScience 23, 101033 (2020).

61. Morton, G.J., Meek, T.H. & Schwartz, M.W. Neurobiology of food intake in health and disease. Nat Rev Neurosci 15, 367–378 (2014).

62. Chometton, S., et al. A premammillary lateral hypothalamic nuclear complex responds to hedonic but not aversive tastes in the male rat. Brain Struct Funct 221, 2183–2208 (2016).

63. Ding, W., Weltzien, H., Peters, C. & Klein, R. Nausea-induced suppression of feeding is mediated by central amygdala Dlk1-expressing neurons. Cell Rep 43, 113990 (2024).

64. Madisen, L., et al. A robust and high-throughput Cre reporting and characterization system for the whole mouse brain. Nature Neuroscience 13, 133–140 (2010).

65. Lao, Z., Raju, G.P., Bai, C.B. & Joyner, A.L. MASTR: a technique for mosaic mutant analysis with spatial and temporal control of recombination using conditional floxed alleles in mice. Cell Rep 2, 386–396 (2012).

66. Chaveroche, M.K., Ghigo, J.M. & d’Enfert, C. A rapid method for efficient gene replacement in the filamentous fungus Aspergillus nidulans. Nucleic Acids Res 28, E97 (2000).

67. Depaepe, V., et al. Ephrin signalling controls brain size by regulating apoptosis of neural progenitors. Nature 435, 1244–1250 (2005).

68. Johansson, T., et al. Building a zoo of mice for genetic analyses: a comprehensive protocol for the rapid generation of BAC transgenic mice. Genesis 48, 264–280 (2010).

69. Raymond, C.S. & Soriano, P. High-efficiency FLP and PhiC31 site-specific recombination in mammalian cells. PLoS One 2, e162 (2007).

