## Supplementary materials for "Food and water uptake are regulated by distinct central amygdala circuits revealed using intersectional genetics"

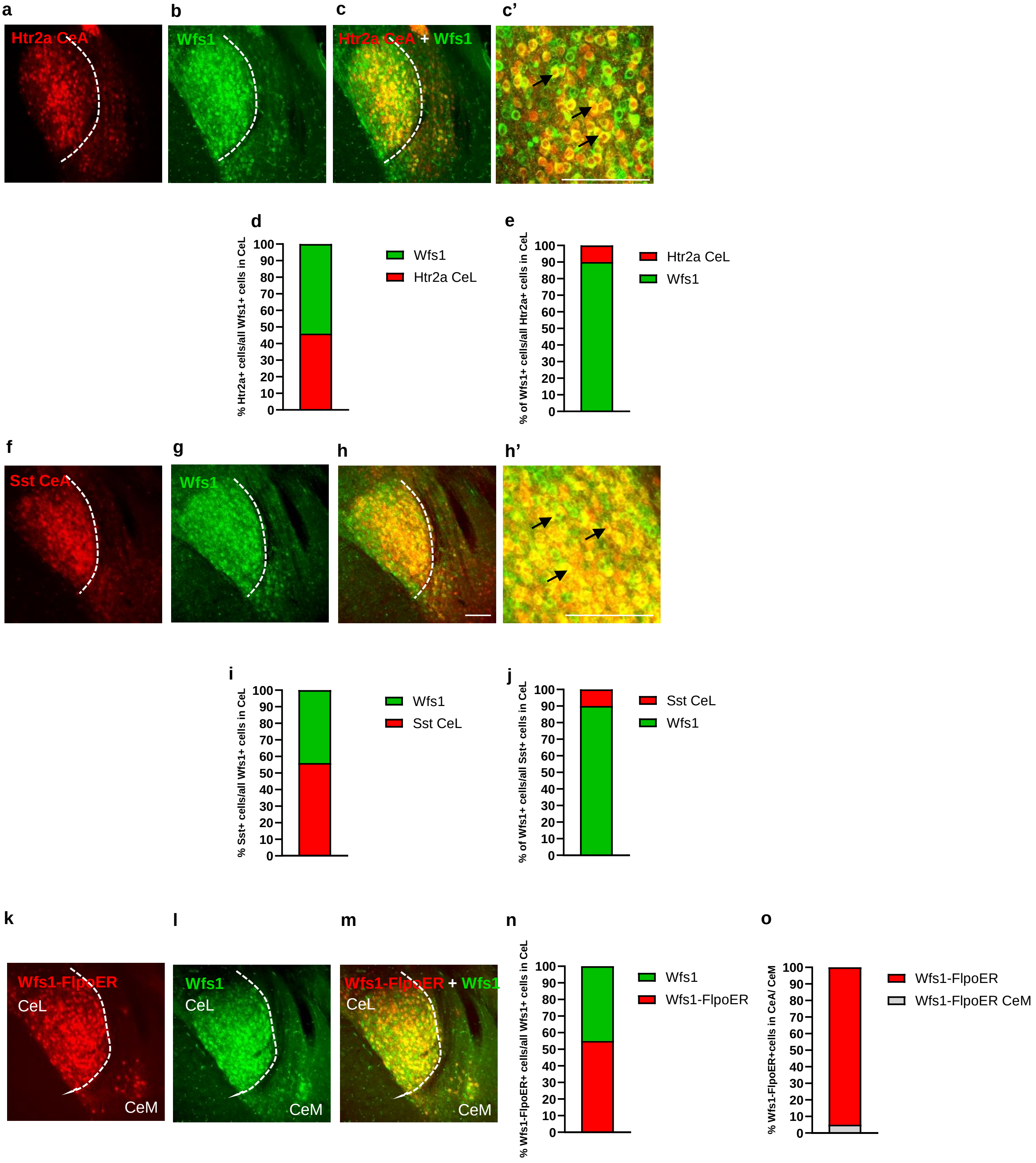


**Extended Data Fig.1**

**Extended Data Figure 1 related to Figure 1. An intersectional strategy to dissect CeL and CeM subpopulations**

**a.** tdTomato fluorescence in adult Htr2a-Cre;Ai9 mice. Stippled white line indicates the border between CeL and CeM.

**b.** Endogenous Wfs1 immunoreactivity in CeL.

**c.** Merge of panels a and b. **c'**. High magnification image of panel c. Black arrows highlight examples of cells double positive for both tdTom and Wfs1.

**d.** Fraction of Htr2a-Cre expressing neurons (46.8±0.3%) among all Wfs1-positive cells in CeL (n = 9 sections from 3 mice).

**e.** Fraction of Wfs1-positive neurons (89.6±1.2%) among all Htr2a-Cre expressing cells in CeL (n = 9 sections from 3 mice).

**f.** Expression of tdTomato in adult Sst-Cre;Ai9 mice.

**g.** Endogenous Wfs1 immunoreactivity in CeL.

**h.** Merge of panels f and g. **h'**. High magnification image of panel h. Black arrows highlight examples of cells colocalizing of both Sst and Wfs1.

**i.** Fraction of Sst-Cre-expressing cells (56.2±1.4%) among all Wfs1-positive cells in CeL (9 sections/3 mice).

**j.** Fraction of Wfs1-positive neurons (89.8±0.3%) among all Sst-Cre-expressing cells in CeL (9 sections/3 mice).

**k-o.** Validation of FlpoER expression. **k.** The activity of tamoxifen-inducible FlpoER in the CeA of Wfs1-FlpoER;FPDi mice leads to expression of mCherry in CeL.

**l.** Endogenous Wfs1 immunoreactivity in CeL.

**m.** Merge of panels k and l.

**n.** Fraction of cells expressing mCherry (transgenic Wfs1-FlpoER; 55.5±4.8%) among all Wfs1-positive cells in CeL (n=3 mice, 3 sections per brain).

**o.** Fraction of mCherry-positive cells in the CeM (5.8±1.4%) among all mCherry- positive cells in the CeA.

Scale bar: 115 µm.

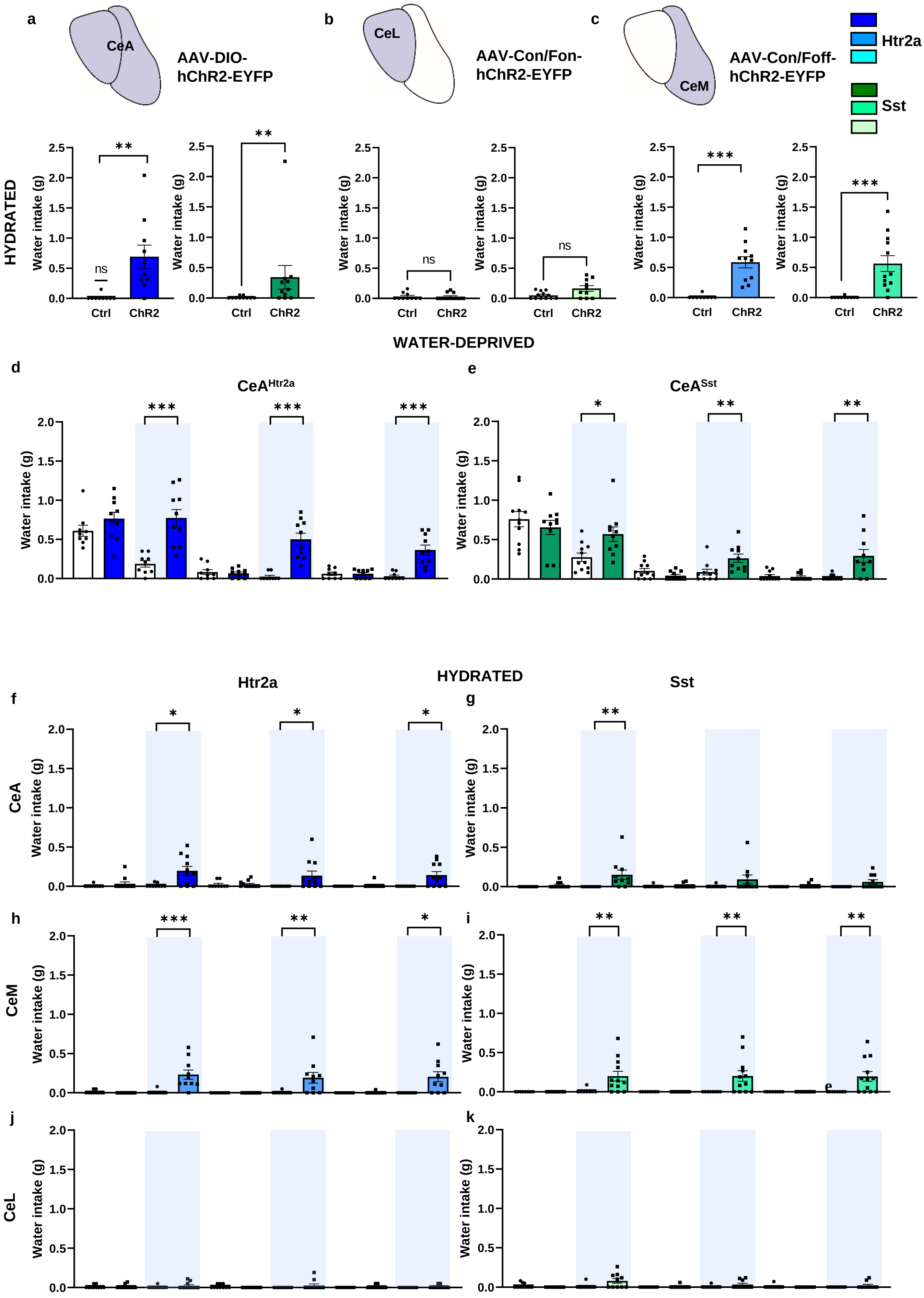


Extended Data Fig.2

**Extended Data Figure 2 related to Figure 2: Water consumption induced by photoactivation of different subpopulations of Htr2a-Cre and Sst-Cre expressing neurons**

**a-c.** Water intake during 30 min by normally hydrated mice during photoactivation of (**a**) the entire CeA populations of Htr2a-Cre (blue) or Sst-Cre expressing neurons (green), (**b**) the respective CeL subpopulations, or (**c**) the respective CeM subpopulations, compared to photoactivated control animals ($CeA^{Htr2a}$: Mann-Whitney U test *p*=0.0002, U=5; $CeA^{Sst}$: Mann-Whitney U test *p*=0.0038, U=18; $CeL^{Htr2a}$: Mann-Whitney U test *p*=0.8939, U=58; $CeL^{Sst}$: Mann-Whitney U test *p*=0.0699, U=33; $CeM^{Htr2a}$: Mann-Whitney U test *p*<0.0001, U=0; $CeM^{Sst}$: Mann-Whitney U test *p*<0.0001, U=5.500).

**d,e.** Water intake during a 10-minute Light OFF/ON behavioral paradigm by water-deprived mice with photoactivated $CeA^{Htr2a}$ (blue) or $CeA^{Sst}$ neurons (green), compared to controls (Htr2a: 10-20 min: unpaired t test *p*=0.0002, t=4.729. 30-40 min: Mann-Whitney U test *p*<0.0001, U=0. 50-60 min: Mann-Whitney U test *p*<0.0001, U=1.500. Sst: 10-20 min: unpaired t test *p*=0.0103, t=2.847. 30-40 min: Mann-Whitney U test *p*=0.0038, U=15.50. 50-60 min: Mann-Whitney U test *p*=0.0014, U=14).

**f-k.** The same 10-minute Light OFF/ON paradigm was repeated with hydrated animals.

**f,g.** Photoactivation of the entire populations of $CeA^{Htr2a}$ or $CeA^{Sst}$ neurons promoted drinking (Htr2a: 10-20 min: Mann-Whitney U test *p*=0.0107, U=18.50. 30-40 min: Wilcoxon signed rank test *p*=0.0156. 50-60 min: Wilcoxon signed rank test *p*=0.0156. Sst: 10-20 min: Wilcoxon signed rank test *p*=0.0078).

**h,i.** Photoactivation of the $CeM^{Htr2a}$ or $CeM^{Sst}$ subpopulations was sufficient to induce water uptake (Htr2a: 10-20 min Mann-Whitney U test *p*=0.0001, U=5.500. 30-40 min: Mann-Whitney U test *p*=0.0031, U=16.50. 50-60 min: Wilcoxon signed rank test *p*=0.0156. Sst: 10-20 min Mann-Whitney U test *p*=0.0067, U=15.50. 30-40 min: Wilcoxon signed rank test *p*=0.0078. 50-60 min: Wilcoxon signed rank test *p*=0.0078).

**j,k**. No effect was observed after photoactivation of the $CeL^{Htr2a}$ or $CeL^{Sst}$ subpopulations (Htr2a: 10-20, 30-40, 50-60 min: Wilcoxon signed rank test or Mann-Whitney U test *p*>0.05. Sst: 10-20, 30-40, 50-60 min: Wilcoxon signed rank test or Mann-Whitney U test *p*>0.05). Value = Mean ± SEM.

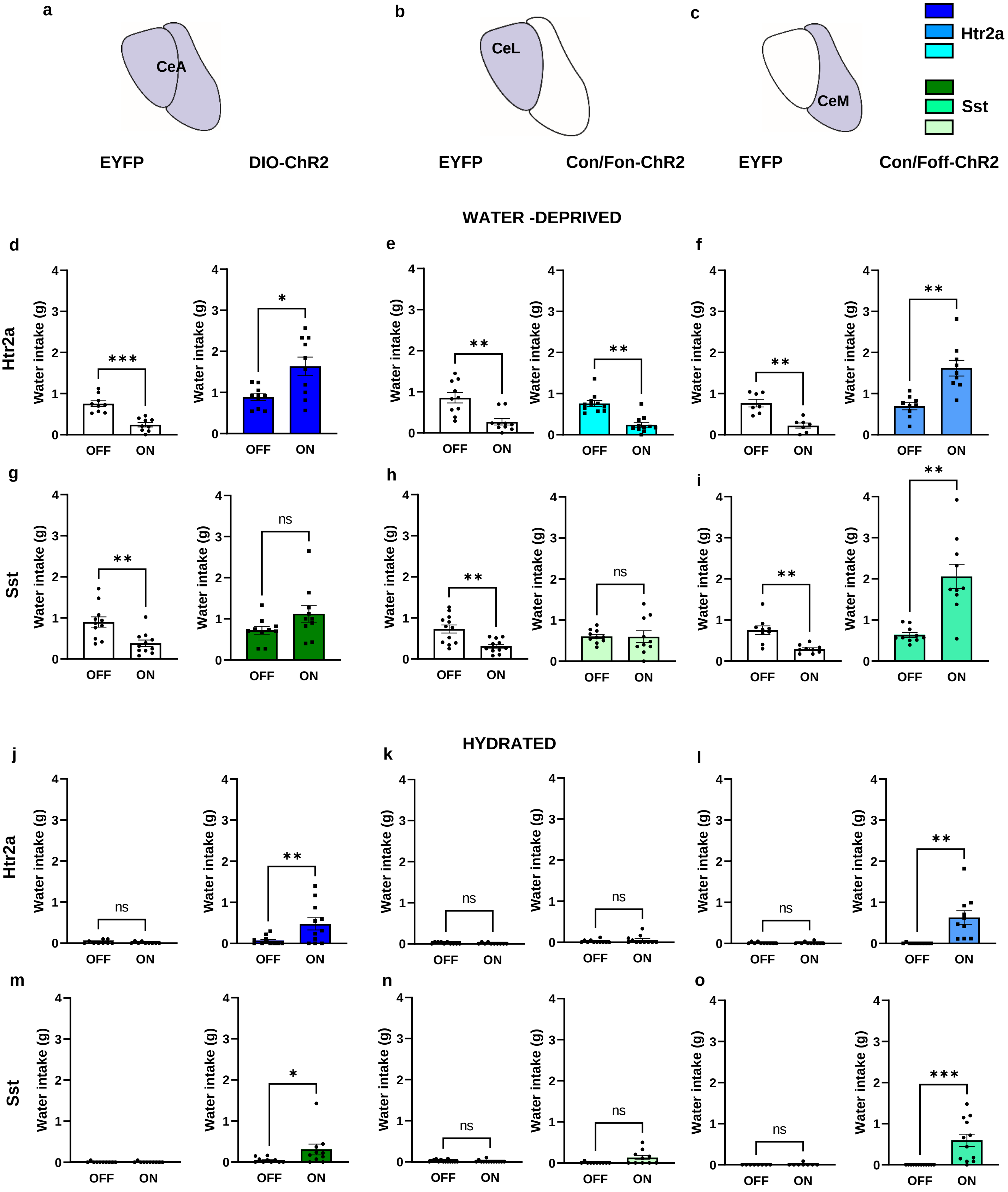


Extended Data Fig.3

**Extended Data Figure 3 related to Figure 2: Comparisons of water consumption between photostimulated and unstimulated ChR2-expressing mice and controls**

**a-c.** Schemes of CeA subregional expression of INTRSECT viruses.

**d-i.** Total amount of water consumed by water-deprived animals during a 10-min light-OFF epoch followed by a 10-min light-ON epoch.

**d.** Photostimulation of the entire population of $CeA^{Htr2a}$ cells promoted water consumption compared to Light OFF periods. The opposite effect was observed in $CeA^{Htr2a}$ controls (Htr2a::ChR2: paired t test *p*=0.0143, t=3.028. Htr2a::EYFP: paired t test *p*=0.0006, t=5.434.).

**e.** No positive effect on water uptake of photostimulating $CeL^{Htr2a}$ or controls (Htr2a::EYFP: Wilcoxon matched-pairs signed rank test *p*=0.0059; Htr2a;ChR2: *p*=0.0029).

**f.** Photostimulation of $CeM^{Htr2a}$ cells increased water uptake compared to light OFF periods. No positive effect of the light in $CeM^{Htr2a}$ controls (Htr2a::EYFP: paired t test *p*=0.0037, t=4.602. Htr2a::ChR2: *p*=0.0059, t=3.720).

**g.** Photostimulation of the entire population of $CeA^{Sst}$ cells did not promote water consumption compared to Light OFF periods. No positive effect of the light in $CeA^{Sst}$ controls (Sst::ChR2: paired t test *p*=0.1054, t=1.800; Sst::EYFP: paired t test *p*=0.0012, t=4.486).

**h.** No positive effect on water uptake of photostimulating $CeL^{Sst}$ or controls (Sst::EYFP: paired t test *p*=0.0018, t=4.079. Sst::ChR2: paired t test *p*=0.9805, t=0.02512).

**i.** Photostimulation of $CeM^{Sst}$ cells increased water uptake compared to light OFF periods. No positive effect of the light in $CeM^{Sst}$ controls (Sst::EYFP: paired t test *p*=0.0017, t=4.625. Sst::ChR2: *p*=0.0014, t=4.568).

**j-o.** Total water intake by hydrated mice during the 10-minutes Light ON/OFF paradigm.

**j.** Photostimulation of the entire population of $CeA^{Htr2a}$ cells promoted water consumption compared to Light OFF periods. No positive effect of the light in $CeA^{Htr2a}$ controls (Htr2a::EYFP: Wilcoxon matched-pairs signed rank test *p*=0.6250; Htr2a::ChR2: *p*=0.0078).

**k.** No positive effect on water uptake of photostimulating $CeL^{Htr2a}$ cells or controls (Htr2a::EYFP: Wilcoxon matched-pairs signed rank test *p*=0.6250; Htr2a::ChR2: *p*=0.1250).

**l.** Photostimulation of $CeM^{Htr2a}$ cells increased water uptake compared to light OFF periods. No positive effect of the light in $CeM^{Htr2a}$ controls (Htr2a::EYFP: Wilcoxon matched-pairs signed rank test *p*>0.9999; Htr2a::ChR2: *p*=0.0020).

**m.** Photostimulation of the entire population of $CeA^{Sst}$ cells did not promote water consumption compared to Light OFF periods. No positive effect of the light in $CeA^{Sst}$ controls (Sst::EYFP: Wilcoxon matched-pairs signed rank test $p>0.9999$. Sst::ChR2: $p=0.0391$).

**n.** No positive effect on water uptake of photostimulating $CeL^{Sst}$ cells or controls (Sst::EYFP: Wilcoxon matched-pairs signed rank test $p=0.5000$. Sst::ChR2: $p=0.0625$).

**o.** Photostimulation of $CeM^{Sst}$ cells increased water uptake compared to light OFF periods. No positive effect of the light in $CeM^{Sst}$ controls (Sst::EYFP: Wilcoxon matched-pairs signed rank test $p>0.9999$; Sst::ChR2: $p=0.0010$). Value = Mean ± SEM.

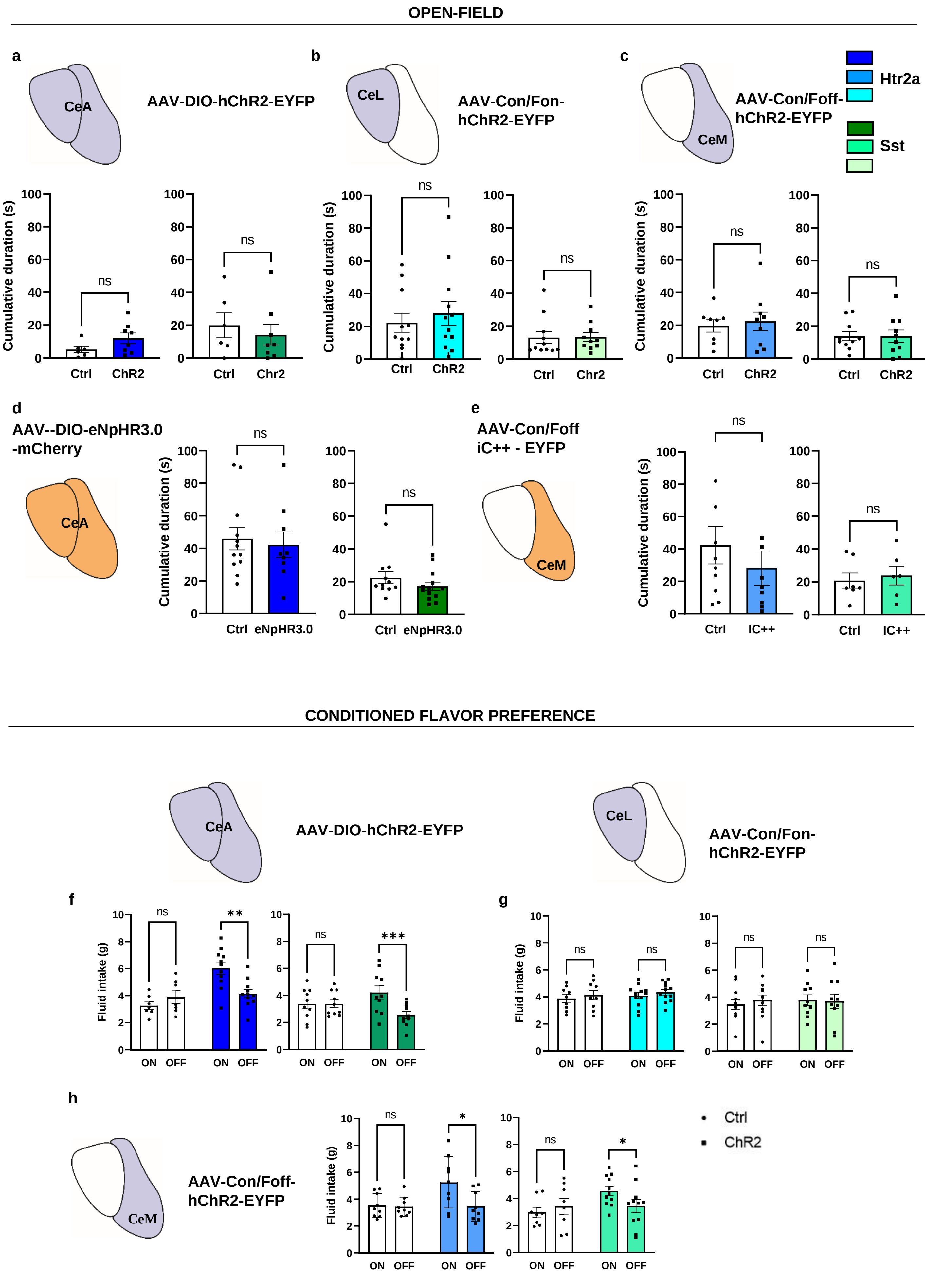


Extended Data Fig.4

**Extended Data Figure 4 related to Figure 4: Effects of optogenetic manipulation of Htr2a and Sst subpopulations in open field and conditioned flavor preference task**

**a.** Cumulative duration in the center-zone during an open-field behavioral test (OF) of mice in which the entire populations of $CeA^{Htr2a}$ (blue) or $CeA^{Sst}$ (green) neurons were photostimulated in comparison to control mice ($CeA^{Htr2a}$: unpaired t test $p$=0.1222, t=1.663. $CeA^{Sst}$: Mann-Whitney U test $p$=0.5987, U=19.50).

**b.** Cumulative duration in the center-zone during OF of mice in which only $CeL^{Htr2a}$ (blue) or $CeL^{Sst}$ (green) neurons were photostimulated in comparison to control mice ($CeL^{Htr2a}$: unpaired t test $p$=0.5541, t=0.6013. $CeL^{Sst}$: Mann-Whitney U test $p$=0.3960, U=42.50.)

**c.** Cumulative duration in the center-zone during OF of mice in which only $CeM^{Htr2a}$ (blue) or $CeM^{Sst}$ (green) neurons were photostimulated in comparison to control mice ($CeM^{Htr2a}$: unpaired t test $p$=0.6934, t=0.4020. $CeM^{Sst}$: $p$=0.9833, t=0.02123).

**d.** Cumulative duration in the center-zone during OF of mice in which the entire populations of $CeA^{Htr2a}$ (blue) or $CeA^{Sst}$ (green) neurons were photoinhibited in comparison to control mice ($CeA^{Htr2a}$: Mann-Whitney U test $p$=0.8480, U=51. $CeA^{Sst}$: $p$=0.1500, U=46).

**e.** Cumulative duration in the center-zone during OF of mice in which only $CeM^{Htr2a}$ (blue) or $CeM^{Sst}$ (green) neurons were photoinhibited in comparison to control mice ($CeM^{Htr2a}$: Mann-Whitney U test $p$=0.2775, U=31. $CeM^{Sst}$: unpaired t test $p$=0.6783, t=0.4260).

**f.** Fluid intake during conditioned flavor preference task by mice in which the entire populations of $CeA^{Htr2a}$ (blue) or $CeA^{Sst}$ (green) neurons were photostimulated in comparison to unstimulated mice (Htr2a: main effect ChR2: Two-way ANOVA, $F$(1,16) =11.94, $p$=0.0033; Bonferroni post-hoc test $p$=0.0014. Sst: main effect Light, Two-way ANOVA, $F$(1,18) =10.30, $p$=0.0049; Bonferroni post-hoc test $p$=0.0005).

**g.** Fluid intake by mice in which only $CeL^{Htr2a}$ (blue) or $CeL^{Sst}$ (green) neurons were photostimulated (Htr2a: Two-way ANOVA $p$>0.05. Sst: Two-way ANOVA $p$>0.05).

**h.** Fluid intake by mice in which only $CeM^{Htr2a}$ (blue) or $CeM^{Sst}$ (green) neurons were photostimulated (Htr2a: main effect Light, Two-way ANOVA, $F$(1,20) =5.888, $p$=0.0248; Bonferroni post-hoc test $p$=0.0014. Sst: main effect LightxChR2, Two-way ANOVA, $F$(1,17) =4.966, $p$=0.0396; Bonferroni post-hoc test $p$=0.0499). Value = Mean ± SEM.

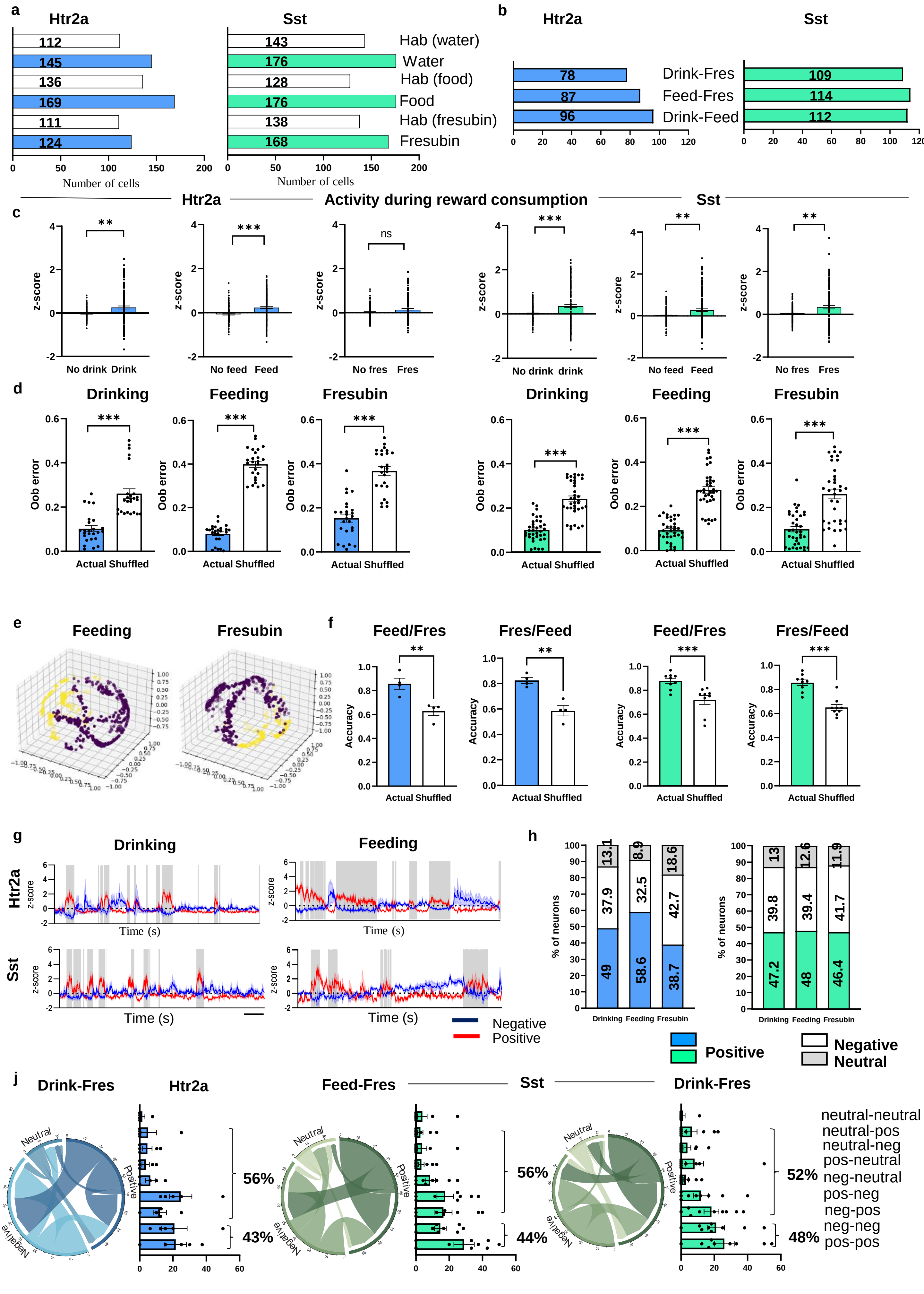


Extended Data Fig.5

**Extended Data Figure 5 related to Figure 5. Calcium responses of CeM$^{Htr2a}$ and CeM$^{Sst}$ neurons to different rewarding stimuli**

**a.** Total numbers of recorded CeM$^{Htr2a}$ and CeM$^{Sst}$ cells in the 10-min habituation and the 10-min reward exposure (water, food or fresubin).

**b.** Numbers of CeM$^{Htr2a}$ and CeM$^{Sst}$ neurons longitudinally detected during drinking-feeding, feeding-Fresubin or drinking-Fresubin behavioral paradigms.

**c.** Average z-score comparisons of the general activity of CeM$^{Htr2a}$ and CeM$^{Sst}$ during habituation and 10-minute exposure to reward (Mann-Whitney U test. CeM$^{Htr2a}$: water, $p$=0.1771, U=7322; food, $p$=0.0098, U=9440; Fresubin, $p$=0.0001, U=4901. CeM$^{Sst}$: water, $p$<0.0001, U=8130; food, $p$<0.0001, U=7320; Fresubin $p$<0.0001, U=8119).

**d.** Out of bag error of Random Forest for Htr2a and Sst embeddings (Htr2a Drinking, Feeding, Fresubin: Wilcoxon matched-pairs signed rank test $p$<0.0001. Sst Drinking, Feeding, Fresubin: Wilcoxon matched-pairs signed rank test $p$<0.0001).

**e.** Representative images of CEBRA generated embeddings of Htr2a neurons longitudinally detected in feeding-Fresubin.

**f.** Accuracy of behavioral decoding through Random Forest, employing CEBRA embeddings of Htr2a (in blue) and Sst (in green) neurons longitudinally detected during feeding and Fresubin consumption. The embeddings derived from feeding behavior are used for predicting Fresubin and vice versa (Htr2a: Feed/Fres: paired t test $p$=0.0042, t=7.891. Fres/Feed: paired t test $p$=0.0043, t=7.838. Sst: Feed/Fres: paired t test $p$=0.0004, t=5.408. Fres/Feed: paired t test $p$<0.0001, t=7.104).

**g.** Representative average activity traces of CeM$^{Htr2a}$ and CeM$^{Sst}$ neurons recorded from one mouse each during 10-minute exposure to water or food. In red the positive and in blue the negative correlated cells (average activity ± SEM). In grey the consummatory bouts.

**h.** Graphs showing the percentages of neurons positively or negatively correlated (blue for CeM$^{Htr2a}$, green for CeM$^{Sst}$) to drinking, feeding and Fresubin consumption.

**j.** Chord diagrams and bar graphs depicting stable or unstable correlations of the same CeM$^{Htr2a}$ and CeM$^{Sst}$ neurons from drinking to Fresubin and from feeding to Fresubin. (Htr2a drinking-Fresubin stable v.s. unstable: Mann-Whitney U test $p$=0.1587, U=4.500. Sst feeding-Fresubin stable v.s. unstable: unpaired t test $p$=0.3176, t=1.032. Sst drinking-Fresubin stable v.s. unstable: unpaired t test $p$=0.6533, t=0.4577). Value = Mean ± SEM.

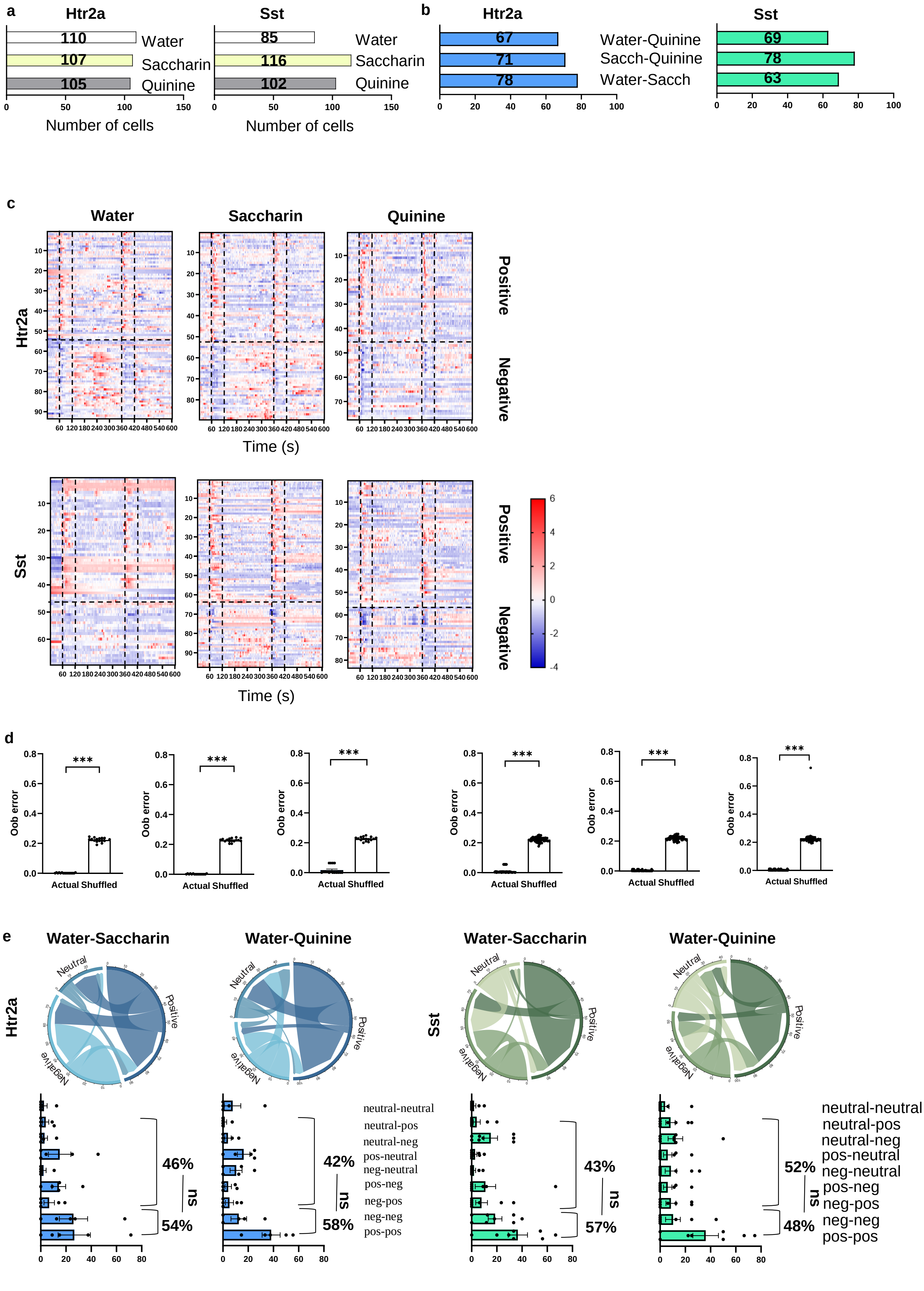


Extended Data Fig.6

**Extended Data Figure 6 related to Figure 6. Calcium responses of $CeM^{Htr2a}$ and $CeM^{Sst}$ cells to different salient stimuli**

**a.** Total numbers of $CeM^{Htr2a}$ and $CeM^{Sst}$ cells recorded during the 10-min water, saccharin and quinine sessions.

**b.** Numbers of $CeM^{Htr2a}$ and $CeM^{Sst}$ neurons longitudinally detected during water-saccharin, saccharin-quinine and water-quinine sessions.

**c.** Heatmap visualization of the activities of $CeM^{Htr2a}$ and $CeM^{Sst}$ neurons having positive or negative correlations with water, saccharin and quinine.

**d.** Out of bag error of Random Forest for Htr2a and Sst embeddings (Htr2a water,saccharin,quinine: Wilcoxon matched-pairs signed rank test $p$<0.0001. Sst water,saccharin,quinine: Wilcoxon matched-pairs signed rank test $p$<0.0001).

**e.** Chord diagrams and bar graphs depicting the stable or unstable correlation (generalizer v.s. specializers) of the same $CeM^{Htr2a}$ and $CeM^{Sst}$ neurons from water to saccharin and from water to quinine. (Htr2a: water-saccharin generalizer v.s. specializers: unpaired t test $p$=0.4554, t=0.7844. Water-quinine generalizer v.s. specializers: unpaired t test $p$=0.3546, t=0.9826. Sst: water-saccharin generalizer v.s. specializers: Mann-Whitney U test $p$=0.0859, U=15. Water-quinine generalizer v.s. specializers: unpaired t test $p$=0.7155, t=0.3719). Value = Mean ± SEM.

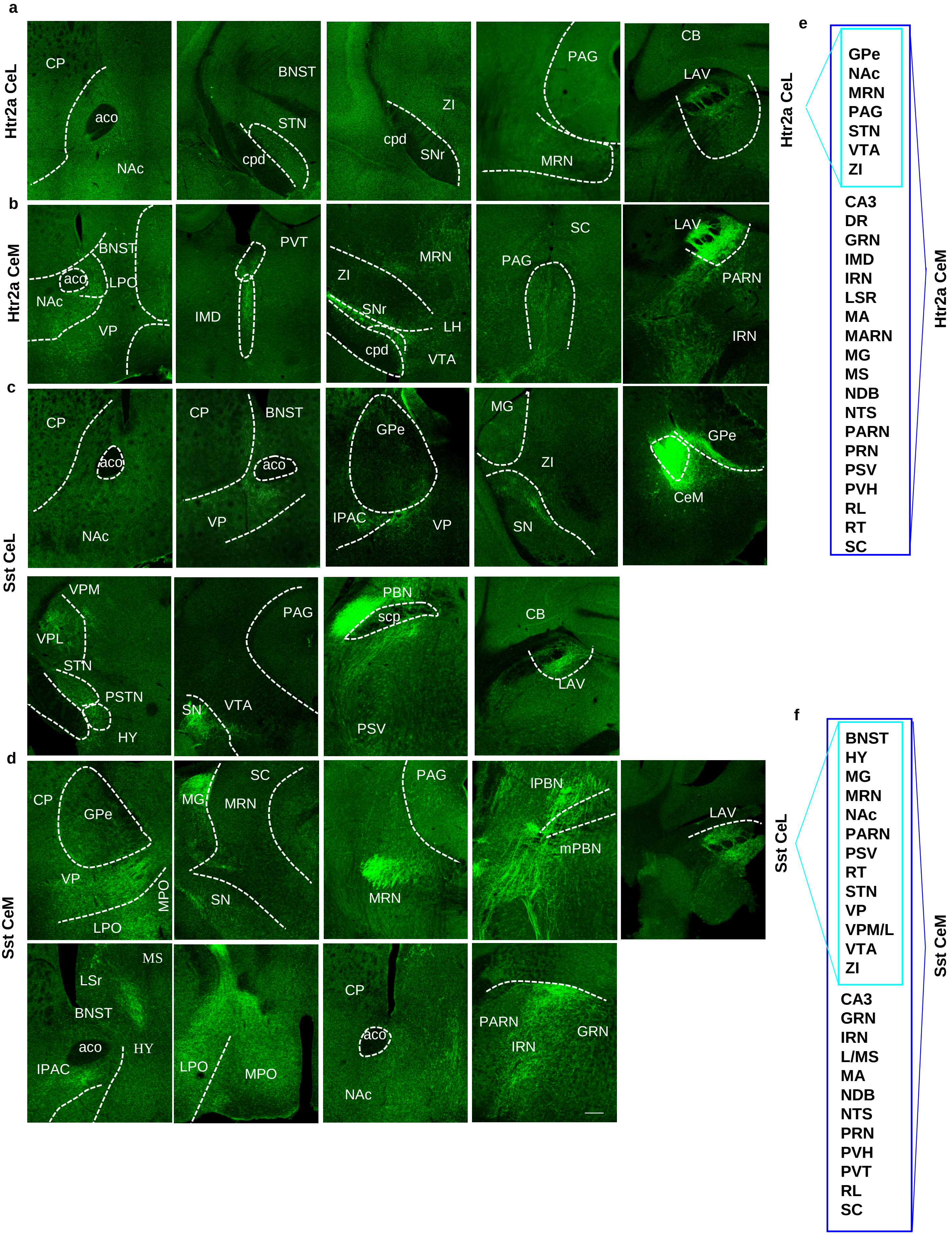


**Extended Data Fig.7**

**Extended Data Figure 7 related to Figure 8. Long range projections of CeL and CeM subpopulations**

**a,b.** Images displaying brain areas receiving projections from CeL/CeM$^{Htr2a}$ neurons.

**c,d.** Images displaying brain areas receiving projections from CeL/CeM$^{Sst}$ neurons. In these experiments Htr2a-Cre;Wfs1-FlpoER or Sst-Cre;Wfs1-FlpoER mice were injected either with a Con/Fon-EYFP or with a Con/Foff-EYFP virus.

**e,f.** Schematic representation of the diverse projections of Htr2a or Sst neurons. BNST: bed nucleus of the stria terminalis. DR: dorsal raphe. GPe: globus pallidus. GRN: gigantocellular reticular nucleus. HY: hypothalamus. IMD: intermediodorsal nucleus of the thalamus. IRN: intermediate reticular nucleus. L/M/Sr: lateral/medial septal nucleus, (rostral part). MA: magnocellular nucleus. MARN: magnocellular reticular nucleus. MG: medial geniculate complex. MPO: medial preoptic area. MRN: midbrain reticular. MS: medial septal nucleus. NAc: nucleus accumbens. NDB: diagonal band nucleus. NTS: nucleus of the solitary tract. PAG: periaqueductal gray. PARN: parvicellular reticular nucleus. PRN: pontine reticular nucleus. PSV: principal sensory nucleus of the trigeminal. PVH: paraventricular nucleus of the hypothalamus. PVT: paraventricular nucleus of the thalamus. RL: rostral linear nucleus raphe. RT: reticular nucleus of the thalamus. SC: superior colliculus. STN: subthalamic nucleus. VP: ventral pallidum. VPM/L: ventral posteromedial/lateral nucleus of the thalamus. VTA: ventral tegmental area. ZI: zona incerta. Scale bar: 139 µm.

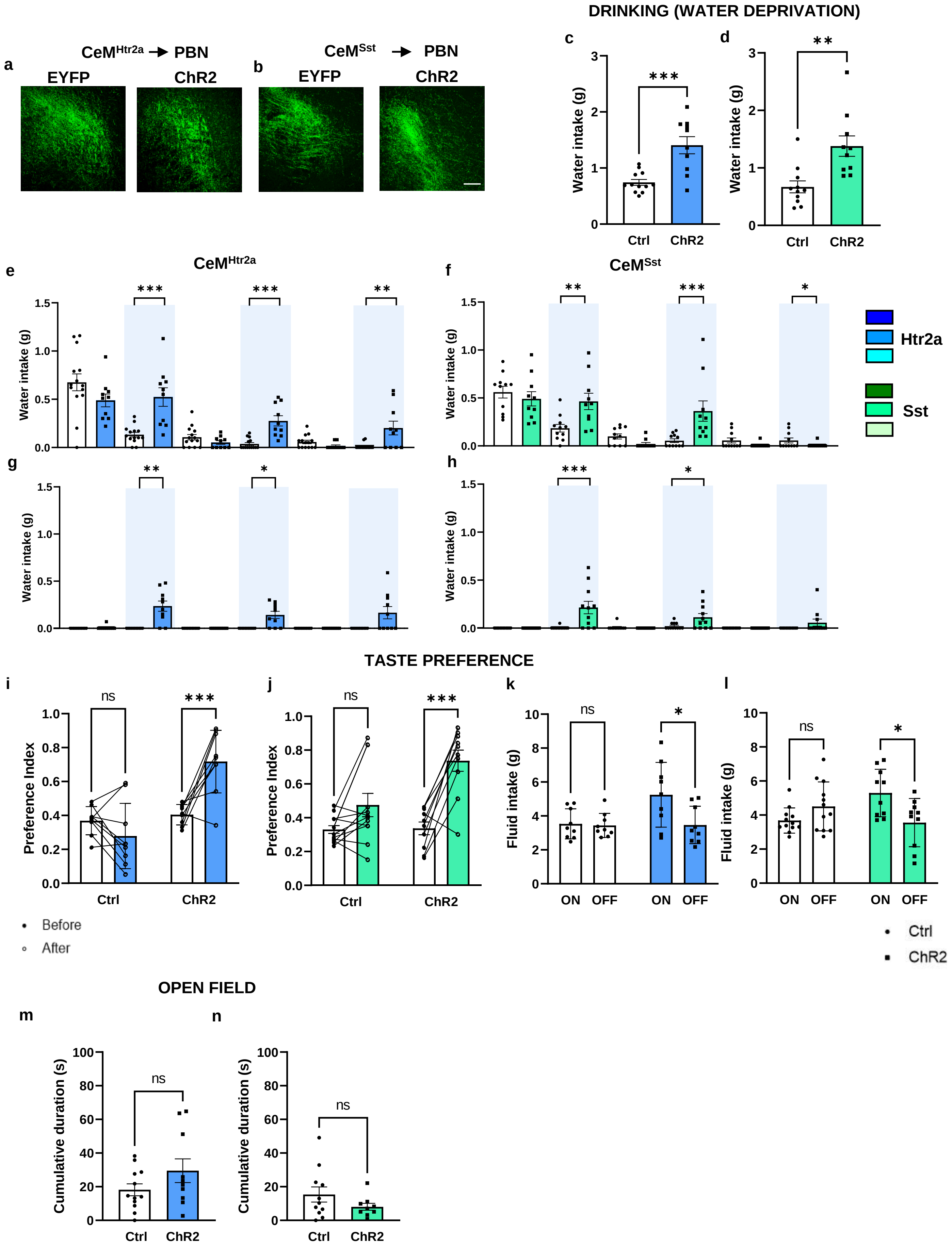


Extended Data Fig.8

**Extended Data Figure 8 related to Figure 8. CeM projections to PBN drive appetitive behaviors**

**a,b.** Representative images displaying the viral expression of ChR2 and EYFP of $CeM^{Htr2a}$ **(e)** and $CeM^{Sst}$ **(f)** projections to PBN.

**c,d.** Activation of $CeM^{Htr2a}$ **(c)** and $CeM^{Sst}$ **(d)** terminals to PBN increases water intake in water deprived animals (Htr2a: unpaired t test *p*=0.0003, t=4.410, Sst: unpaired t test *p*=0.0022, t=3.531).

**e-f.** Htr2a/Sst-Cre;Wfs1-FlpoER mice, water deprived, expressing ChR2 or EYFP in CeM neurons and photostimulated in the PBN, were exposed to water in a 10 min OFF/ON behavioral paradigm. The test comprised three cycles of 10 minutes without light followed by 10 minutes with light.

**e.** Activation of $CeM^{Htr2a}$ PBN terminals significantly increased drinking behavior (10-20 min: unpaired t test *p*=0.0002, t=4.544, 30-40 min: Mann-Whitney U test *p*<0.0001, U=6, 50-60 min Mann-Whitney U test *p*=0.0074, U=32).

**f.** Activation of $CeM^{Sst}$::ChR2 terminals to the PBN, in the same 10 OFF/ON behavioral paradigm, promoted drinking (10-20 min: unpaired t test *p*=0.0065, t=3.057, 30-40 min: Mann-Whitney U test *p*=0.0003, U=8, 50-60 min: Mann-Whitney U test *p*=0.0136, U=23)

**g-h.** The same experiment was repeated with water-satiated animals and showed that while the control group did not consume much water, photostimulation of $CeM^{Htr2a/Sst}$::ChR2 projections to PBN was sufficient to promote water uptake (Htr2a. 10-20 min: Wilcoxon signed rank test *p*=0.0078, 30-40 min Wilcoxon signed rank test *p*=0.0156) **(g),** (Sst. 10-20 min: Mann-Whitney U test *p*=0.0009, U=20, 30-40 min: Mann-Whitney U test *p*=0.0492, U=36) **(h)**.

**i.** Optogenetic activation of $CeM^{Htr2a}$ terminals to PBN changed the taste preference for the ChR2 expressing mice but not for the control group (main effect ChR2: Two-way ANOVA, *F*(1,16) =22.96, *p*=0.0002; Bonferroni post-hoc test *p*=0.0004).

**j.** Activation of the $CeM^{Sst}$ terminals to PBN reversed the initial preference of the mice (main effect ChR2: Two-way ANOVA, *F*(1,20) =5.585, *p*=0.0284; Bonferroni post-hoc test *p*<0.0001).

**k,l.** Photostimulation during training induced more consumption of flavored water during the light ON period for $CeM^{Htr2a}$::ChR2 (main effect ChR2: Two-way ANOVA, *F*(1,16) =5.888, *p*=0.0182; Bonferroni post-hoc test *p*=0.0383) **(k)** and $CeM^{Sst}$::ChR2 (main effect LightxChR2: Two-way ANOVA, *F*(1,20) =9.879, *p*=0.0051; Bonferroni post-hoc test *p*=0.0184) **(l)** terminals to PBN.

**m,n.** No effect was found on the time spent in the center-zone during an open-field test following activation of $CeM^{Htr2a}$ **(m)** (unpaired t test *p*=0.1447, t=1.518) or $CeM^{Sst}$ **(n)** (unpaired t test *p*=0.1873, t=1.371) PBN projections.

Value = Mean± SEM. Scale bar: 115 µm.
